# Identification of silkworm hemocyte subsets and analysis of their response to BmNPV infection based on single-cell RNA sequencing

**DOI:** 10.1101/2020.10.18.344127

**Authors:** Min Feng, Junming Xia, Shigang Fei, Xiong Wang, Yaohong Zhou, Pengwei Wang, Luc Swevers, Jingchen Sun

**Affiliations:** Guangdong Provincial Key Laboratory of Agro-animal Genomics and Molecular Breeding, College of Animal Science, South ChinaAgricultural University, Guangzhou 510642, China; Insect Molecular Genetics and Biotechnology, National Centre for Scientific Research Demokritos, Institute of Biosciences and Applications, Athens 15310, Greece

**Keywords:** scRNA-seq, *Bombyx mori*, Hemocytes, BmNPV, Lepidopteron

## Abstract

A wide range of hemocyte types exist in insects but a full definition of the different subclasses is not yet established. The current knowledge of the classification of silkworm hemocytes mainly comes from morphology rather than specific markers, so our understanding of the detailed classification, hemocyte lineage and functions of silkworm hemocytes is very incomplete. *Bombyx mori* nucleopolyhedrovirus (BmNPV) is a representative member of the baculoviruses, which are a major pathogens that specifically infects silkworms and cause serious loss in sericulture industry. Here, we performed single-cell RNA sequencing (scRNA-seq) of silkworm hemocytes in BmNPV and mock-infected larvae to comprehensively identify silkworm hemocyte subsets and determined specific molecular and cellular characteristics in each hemocyte subset before and after viral infection. A total of 19 cell clusters and their potential marker genes were identified in silkworm hemocytes. Among these hemocyte clusters, clusters 0, 1, 2, 5 and 9 might be granulocytes (GR); clusters 14 and 17 were predicted as plasmatocytes (PL); cluster 18 was tentatively identified as spherulocytes (SP); and clusters 7 and 11 could possibly correspond to oenocytoids (OE). In addition, all of the hemocyte clusters were infected by BmNPV and some infected cells carried high viral-load in silkworm larvae at 3 day post infection (dpi). Interestingly, BmNPV infection can cause severe and diverse changes in gene expression in hemocytes. Cells belonging to the infection group mainly located at the early stage of the pseudotime trajectories. Furthermore, we found that BmNPV infection suppresses the immune response in the major hemocyte types. In summary, our scRNA-seq analysis revealed the diversity of silkworm hemocytes and provided a rich resource of gene expression profiles for a systems-level understanding of their functions in the uninfected condition and as a response to BmNPV.

## Introduction

The domesticated silkworm, *Bombyx mori*, the only truly domesticated insect,has been used as a model system for lepidopterans for many decades (Xia et al., 2014, Goldsmith et al., 2005). Since ancient times, silkworm breeding has been an important part of the agricultural economy. Starting at the beginning of the 19th century, the silkworm became a model for scientific discovery in physiology, genetics and microbiology and remained in that position for long periods during which large paradigm changes took place in our understanding of biology (Goldsmith et al., 2005). Even to this day, silkworm still plays roles in some important areas such as sericulture, basic research, biotechnology and silk-based biomaterials.

Insects including the silkworm have an efficient and potent innate immune system to discriminate and respond to invading pathogenic microorganism (Sheehan et al., 2020). The innate immune system of insects is divided into humoral defenses that include the production of soluble effector molecules (like antimicrobial peptides) and cellular defenses like phagocytosis and encapsulation that are mediated by hemocytes (Strand, 2008). A wide range of hemocytes exists in insects but a full definition of the different subclasses is not yet established (Sheehan et al., 2020). In Lepidopteran insects such as the silkworm, hemocytes are generally classified into five major subsets including prohemocytes, plasmatocytes, granulocytes, spherulocytes, and oenocytoids based on morphology and function (Lavine and Strand, 2002, Strand, 2008, Ribeiro and Brehelin, 2006). Generally, prohemocytes are considered as multipotent progenitor cells or stem cells giving rise to the other subsets (Yamashita and Iwabuchi, 2001). Plasmatocytes and granulocytes are the only hemocyte subsets capable of adhering to foreign surfaces, and together usually comprise more than 50% of the hemocytes in circulation in the larval stages (Lavine and Strand, 2002). Furthermore, plasmatocytes and granulocytes are involved in most cellular defense responses (Strand, 2008, Lavine and Strand, 2002). Oenocytoids are in rich in polyphenol oxidase and mainly participate in melanization, whereas the functions of spherulocytes are unknown (Nakahara et al., 2010, Nakahara et al., 2009).

Because our current knowledge of the classification of silkworm hemocytes mainly comes from morphology and cytochemistry rather than from specific genetic markers, our detail understanding of hemocyte lineages and functions is very limited. Importantly, if hemocyte subsets cannot be distinguished well, the molecular responses of each hemocyte subset during the invasion of exogenous pathogenic microorganisms remain inaccessible. In addition, although some research teams have studied the hematopoietic lineage of the silkworm (Nakahara et al., 2010, Yamashita and Iwabuchi, 2001), still little is known about silkworm hemocyte lineage trajectories starting from precursor cells or the existence of putative intermediate states during the differentiation process towards mature cell types. Hence, it is important and interesting to thoroughly characterize the molecular signatures and responses of all the precursor and mature hemocyte subtypes during homeostasis and after exposure to pathogenic microorganisms.

Baculoviruses are a very diverse group of invertebrate-specific DNA viruses with genome sizes varying from about 80 to over 180 kb, that encode between 90 and 180 genes, which bring harm to economic insects, but also have been harnessed for biotechnological applications such as insect pest control and the expression of heterologous proteins (Rohrmann, 2019). *Bombyx mori* nucleopolyhedrovirus (BmNPV), a representative member of baculoviruses, is a major pathogen that specifically infects silkworms and causes serious losses in the sericulture industry (Yu et al., 2017). However, no effective measures are currently available to prevent BmNPV infection. With respect to sericulture, many silkworm strains have very little resistance to BmNPV infection which can lead to widespread mortality and population collapse. The general high virulence of baculovirus infections has also been exploited for their successful application as biocontrol agents to kill pests. However, although methods such as bulk RNA-seq and proteomics have been used in recent years to try to clarify the interaction between baculovirus and the host (Yu et al., 2017, Zhang et al., 2020c, Zhang et al., 2020b, Li et al., 2016, Feng et al., 2020, Hu et al., 2018), major gaps remain regarding the host response to the virus, especially at the level of cellular immunity. Hence, the elucidation of the response of different hemocyte subsets to BmNPV infection is a powerful approach to explore the insect cellular immune response to baculovirus infection.

Single-cell RNA sequencing (scRNA-seq) has already been applied to investigate the immune response under the conditions of virus infection (Zhang et al., 2020a, Steuerman et al., 2018). In particular, scRNA-seq technology is widely used to characterize the responses of various cell types to severe acute respiratory syndrome corona virus 2 (SARS-CoV-2) infection (Chua et al., 2020, Zhu et al., 2020, Wilk et al., 2020). In *Drosophila*, one of the major model organisms, the cell lineage characteristics in many tissues have been identified using scRNA-seq, such as the midgut (Hung et al., 2020), blood cells or hemocytes (Tattikota et al., 2020, Cattenoz et al., 2020), ovary (Jevitt et al., 2020, Slaidina et al., 2020), and brain (Davie et al., 2018). However, the application of scRNA-seq technology to determine cell subsets and cell lineages is rare in other insects. Moreover, to our knowledge, scRNA-seq technology has not been applied yet to study the responses of specific insect cell (sub)types to viral infections.

Here, we have performed scRNA-seq on silkworm hemocytes in BmNPV-infected larvae and uninfected larvae to comprehensively identify hemocyte subsets and characterize their specific molecular and cellular responses after viral infection.

## Results

### Single-cell transcriptomics identifies 19 distinct cell clusters in the hemolymph collected from BmNPV-infected and control silkworms

We used the 10x Genomics platforms to perform 3’ scRNA-seq on pooled hemocytes collected from BmNPV-infected silkworm and PBS-treated controls (Figure 1A). Details on the statistics of scRNA-seq are summarized in Supplementary file 1. A total of 22,286 cells (BmNPV: 9,114; Control: 13,172) were profiled and 19 distinct clusters were obtained that can be visualized using t-SNE (Figure 1B, 1C). From cluster 0 (3204 cells, 14.38%) to cluster 18 (47 cells, 0.21%), the number of cells gradually decreases (Figure 1C). However, no known marker genes exist that can be used to specify different types of silkworm hemocytes, in contrast to the robust classification of hemocyte subgroups in *Drosophila* (Tattikota et al., 2020, Cattenoz et al., 2020). Next, we identified the up-regulated DEGs of each cluster and analyzed the enrichment of specific groups of genes involved in distinct cellular processes. Silkworm hemocyte clusters were detected with a large number of up-regulated DEGs, especially cluster 14 (2529 DEGs), 9 (2456), 5 (2062), 2 (2005), 18 (1492), 17 (1458), 0 (1314), 11 (682), 1 (585) and 7 (248) (Figure 1D; Supplementary file 2). It is worth noting that no up-regulated genes were identified in cluster 4 (Figure 1D). The expression levels and the percentage of cells expressing the top five genes in each cluster are shown in a dot plot (Figure 1E; Supplementary file 3). These genes need to be confirmed in future research and are proposed to be used as marker genes for each cluster of silkworm hemocytes (Figure 1E). Through GO and KEGG analysis on all up-regulated DEGs in each cluster, we found that genes involved in immune-related host response processes are enriched in cluster 0, 1, 2, 5, 7, 9, 11, 14, 17, and 18 (Figure 1F, G). We speculate that these hemocyte clusters are the main effectors of silkworm that respond to external stimuli. However, the clusters with the highest numbers of DEGs and enrichment in immune-related host response processes correspond to the clusters that are predominant in control hemocytes (clusters 0, 1, 2, 3, 7, 9, 11, 14, 17 and 18) (see further below).

**Figure 1.**
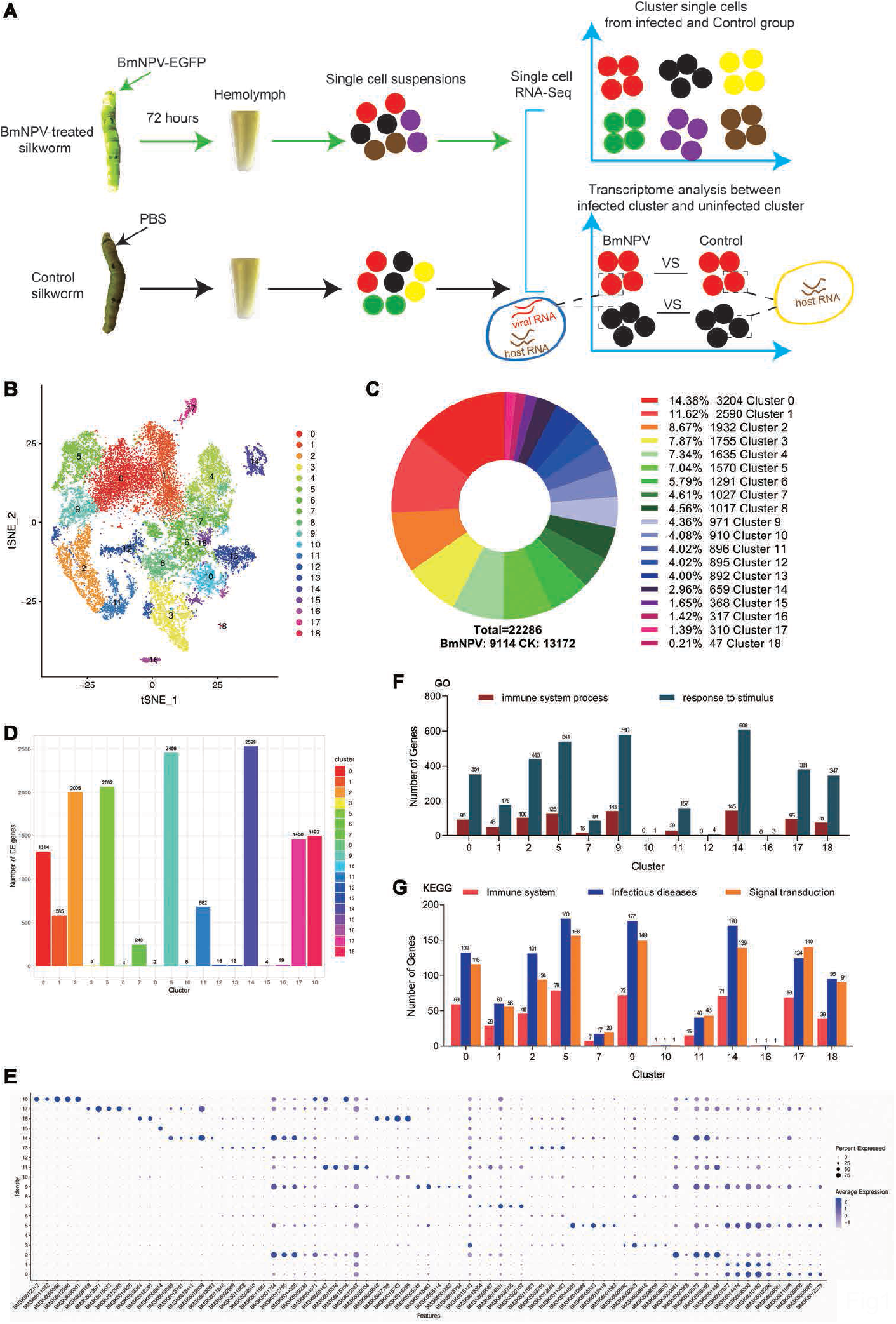
Single-cell profiling of cell populations in silkworm hemolymph. **(A)** Schematic illustration of the experimental workflow. Twenty larvae for each group were treated with BmNPV or PBS and sacrificed at 72 hours post treatment. Hemolymph of the twenty larvae in each group were pooled as one sample for single-cell sequencing (scRNA-seq) analysis. In each single cell, mRNAs of host and virus were simultaneously measured, allowing comparison of the transcriptome profile in each cell cluster under the infection or control condition. **(B)** t-Distributed Stochastic Neighbor Embedding (t-SNE) projection representing the 19 clusters of cells identified in the silkworm hemocyte pools (unified set of control and BmNPV infection samples). **(C)** Number of cells in each cluster and their proportional distribution in the total hemocyte dataset. **(D)** Number of up-regulated differentially expressed genes(DEGs)in each cluster. **(F)** The statistics of genes involved in “immune system process” and “response to stimulus” GO terms as analyzed in each cluster. **(G)** The statistics of genes involved in “Immune system”, “Infectious diseases” and “Signal transduction” KEGG pathways as analyzed in each cluster. **(E)** Top 5 DEGs (x axis) identified in each cluster (y axis). Dot size represents the fraction of cells in the cluster that express the gene; intensity indicates the mean expression (Z-score) in expressing cells, relative to other clusters.

To check that the strategy of combining the hemocytes of the control and BmNPV groups into one set has no effect on the results, the hemocytes of the separate groups of control and BmNPV infection were mapped to the unified set (BmNPV+CK) using cell-specific barcodes. In addition, the hypervariable genes that were identified for cell cluster classification in scRNA-seq were compared between individually analyzed groups and the unified set (all). Figure supplement 1 shows that the agreement between the two clustering approaches is relatively high (Figure supplement 1A, B), especially for the cells in the control group (Figure supplement 1B). Furthermore, irrespective of whether control and BmNPV infection groups were analyzed separately or as a unified set, most of the hypervariable genes to cluster the hemocytes were the same (Figure supplement 1C). These results confirm that the analysis of the unified set of cells from both experimental groups (as used in the remainder of this study) represents a valid approach.

### Immune-related gene expression signatures of each cluster

According to the previously reported silkworm immune-related gene data (including RNAi-related genes) (Tanaka et al., 2008, Kolliopoulou et al., 2015), a total of 105 immune-related genes were screened from cluster 0 to 18 and their expression level was presented as a heat map (Figure 2A; Supplementary file 4). We can observe that most of these immune-related genes were highly expressed in clusters 0, 1, 2, 5, 7, 9, 11, 14, 17 and 18 (Figure 2A). It is well established that RNAi is a major antiviral defense mechanism in insects (Swevers et al., 2018). Besides RNAi, many other innate immune pathways have been proposed to be involved in antiviral defense such as the Imd and Toll pathways, the JAK/STAT pathway, autophagy and apoptosis (Kingsolver et al., 2013, Swevers et al., 2018, Marques and Imler, 2016). The dot plot showed that most of RNAi-related (Figure 2B) and innate immune pathway-related genes (Figure 2C) were highly expressed in clusters 0, 1, 2, 5, 9, 14, 17 and 18. Notably, the antimicrobial peptide (AMP) gene *Cecropin A* was only detected in the cluster 17 and *gloverin 2* was highly expressed in several clusters such as clusters 2, 5, 9, and 14 (Figure 2D). AMP genes of the *Cecropin B* family are highly expressed in hemocyte clusters 5 and 9, especially cluster 9 (Figure 2D). Caspase-8, a molecular switch for apoptosis, necroptosis and pyroptosis (Fritsch et al., 2019), was found to be highly expressed in most of the cells of several clusters such as clusters 0, 5 and 9 (Figure 2E). *Caspase-3*, encodes a key enzyme playing an essential role in both exogenous and endogenous apoptotic pathways (D’Arcy, 2019). We found that the silkworm *Caspase-3* is only highly expressed in cluster 11, while low expression was observed in cluster 2 and expression was restricted to a small number of cells in cluster 7 (Figure 2E). However, *Caspase-1*, one of the critical genes of pyroptosis (Shi et al., 2017), was highly expressed in clusters 14 and 17 (Figure 2E). Death pathways are major defense mechanisms against baculovirus infection (Clem, 2005) and these data therefore indicate that hemocyte clusters 0, 5, 9, 11, 14, and 17 could use this strategy, although different death pathways may be engaged in different clusters.

**Figure 2.**
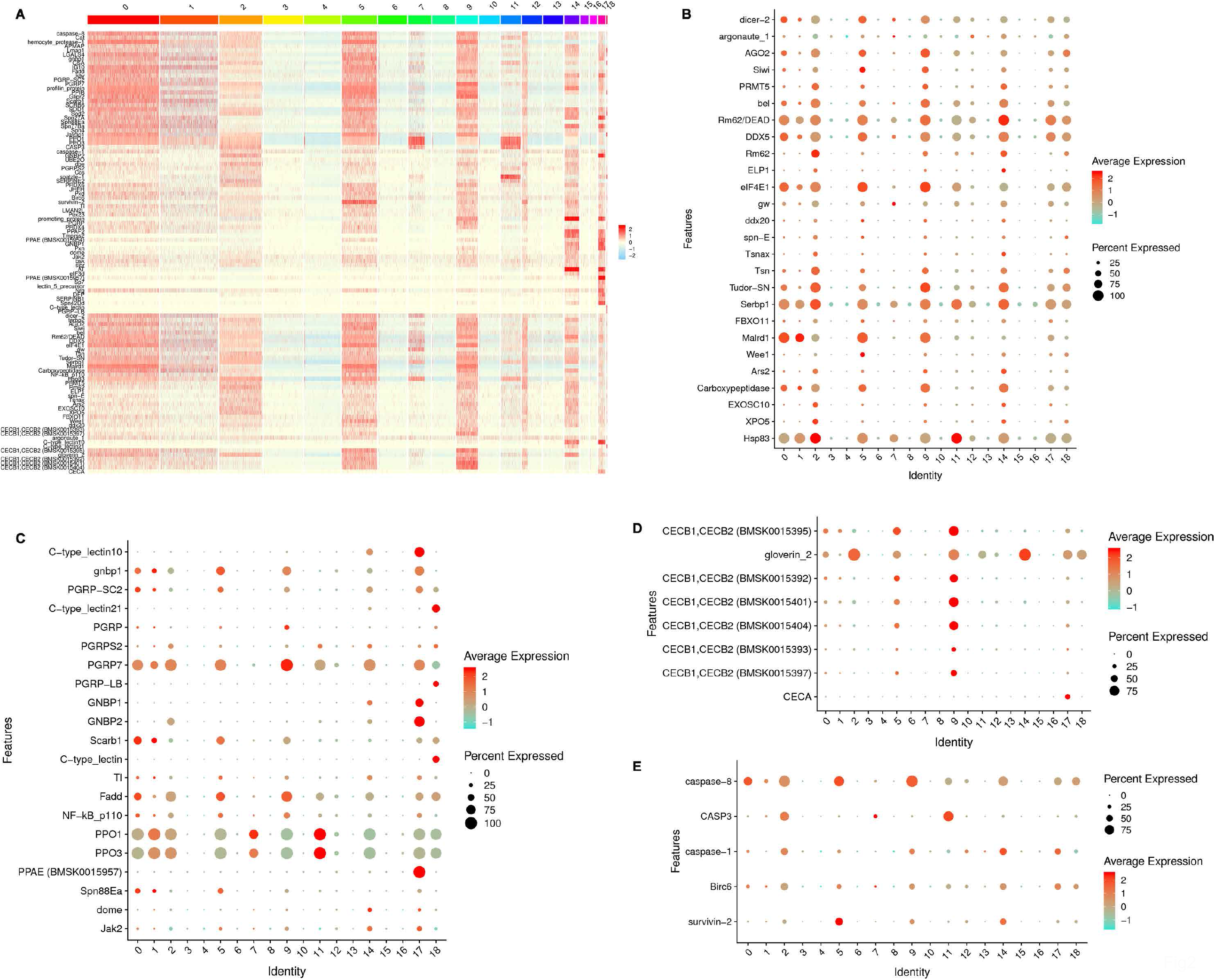
Landscape of expression of immune-related genes in silkworm hemocyte clusters. **(A)** Heatmap showing the normalized expression (Z-score) of all immune-related DEGs in various cells of hemocyte clusters. **(B, C, D, E)** Dot plot representing DEGs belong to “RNAi” (B), “innate immune pathway” (C), “antimicrobial peptide genes” (D) and “cell death” (E) categories in hemocyte clusters based on average expression. Color gradient of the dot represents the expression level, while the size represents the percentage of cells expressing any gene per cluster.

In conclusion, hemocyte clusters 0, 1, 2, 5, 7, 9, 11, 14, 17 and 18 may be involved in the antiviral responses after BmNPV infection.

### Major impact of BmNPV infection on hemocyte identities and physiologies

We further compared the distribution of each silkworm hemocyte cluster in the BmNPV-infected and the control groups. In the uninfected sample, we detected all previously described cell types from cluster 0 to cluster 18 (Figure 3A), although cell numbers in clusters 3, 4, 6, 8, 10, 13, 15 and 16 were much reduced. Interestingly, hemocyte clusters which may be involved in the antiviral response such as clusters 0, 1, 2, 5, 9, 11, 14, and 17 are the main components of uninfected silkworm hemocytes (Figure 3A). On the other hand, only a few cells were detected in these hemocyte clusters (0, 1, 2, 5, 9, 11, 14, and 17) in the BmNPV-infected sample (Figure 3B). Cluster 5 was absent in the BmNPV-infected sample, while only one cell was detected in clusters 0 and 9 (Figure 3B). Correspondingly, few cells of hemocyte clusters 3, 4, 6, 8, 10, 13, 15 and 16 were detected in the control, while a large number of cells were detected in these clusters in the infection group (Figure 3A, B). Comparing the cellular landscapes, we therefore observe that the cells in the BmNPV-infected group and the control group were for a large part distributed in different clusters (Figure 3C). After analyzing the proportion of control and infected cells in each cluster, it was found that hemocyte clusters which may be involved in the antiviral response (0, 1, 2, 5, 9, 11, 14, and 17) showed a striking depletion in the BmNPV-infected sample (Figure 3D). On the other hand, clusters 3, 4, 6, 8, 10, 13, 15 and 16 are the main components of the silkworm hemocytes in the BmNPV-infected group (Figure 3D). Within the clusters that dominated the BmNPV sample, however, few highly expressed DEGs were detected (Figure 1D). We speculate that this reflects baculovirus infection during which virus replication competitively inhibits the expression of host genes. Because very few DEGs were detected in the clusters that appeared in the infection group, gene enrichment analysis could not be performed, preventing characterization of these cell clusters.

**Figure 3.**
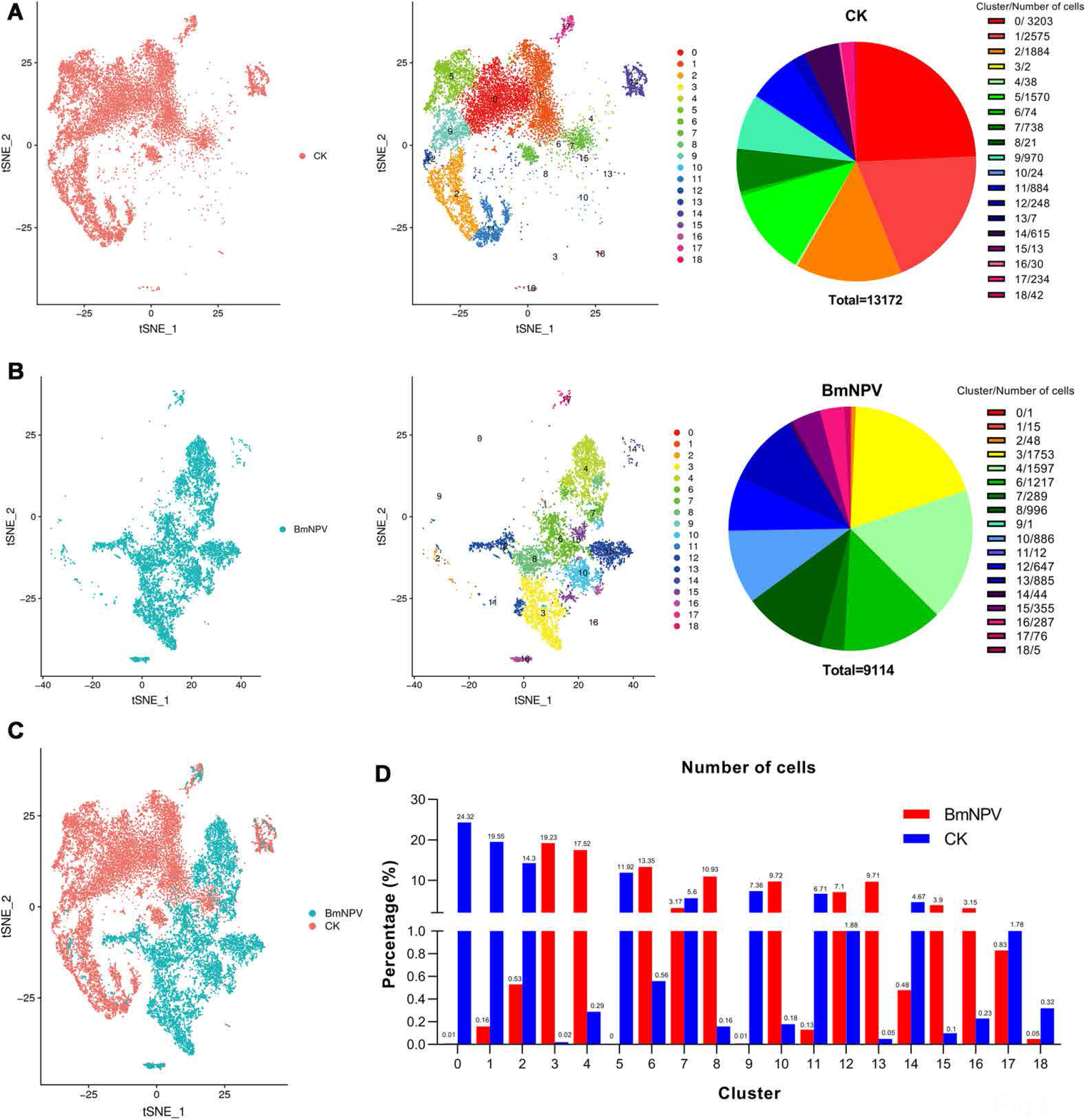
Hemocyte heterogeneity between BmNPV-infected and mock-infected larvae. **(A, B)** t-SNE displaying all identified cell types and their cell number in PBS-treated (CK) (A) and BmNPV-infected larvae (B). **(C)** t-SNE displaying the high heterogeneity in hemocyte clusters between the BmNPV-infected and control groups. **(D)** The distribution of cells in each cluster between the BmNPV-infected and control groups.

### Cell types appearing in the infection group are mainly located at the early stages of the pseudo-time trajectories

A high dimensional graph representation of the data of all hemocyte clusters was constructed and subsequently used for visualization in a low-dimensional UMAP plot (Figure 4A) (Becht et al., 2019). The cell clusters of the infection group and the control showed clear separation in the UMAP plot (Figure 4A). We next sought to investigate the developmental trajectories of different hemocyte clusters in both BmNPV-infected and control samples and the pseudo-times were constructed from all single-cell transcriptomics data using Monocle 2 (Trapnell et al., 2014). Interestingly, the cell clusters highly enriched in the BmNPV-infected sample (3, 4, 6, 8, 10, 13, 15 and 16) (Figure 4A) are mainly located at the early stage of the pseudo-time trajectories (Figure 4B, C; Figure supplement 2). Correspondingly, the hemocyte cluster in the control group (0, 1, 2, 5, 9, 11 and 17) (Figure 4A) are mainly located at the late stage (Figure 4B, C; Figure supplement 2).

**Figure 4.**
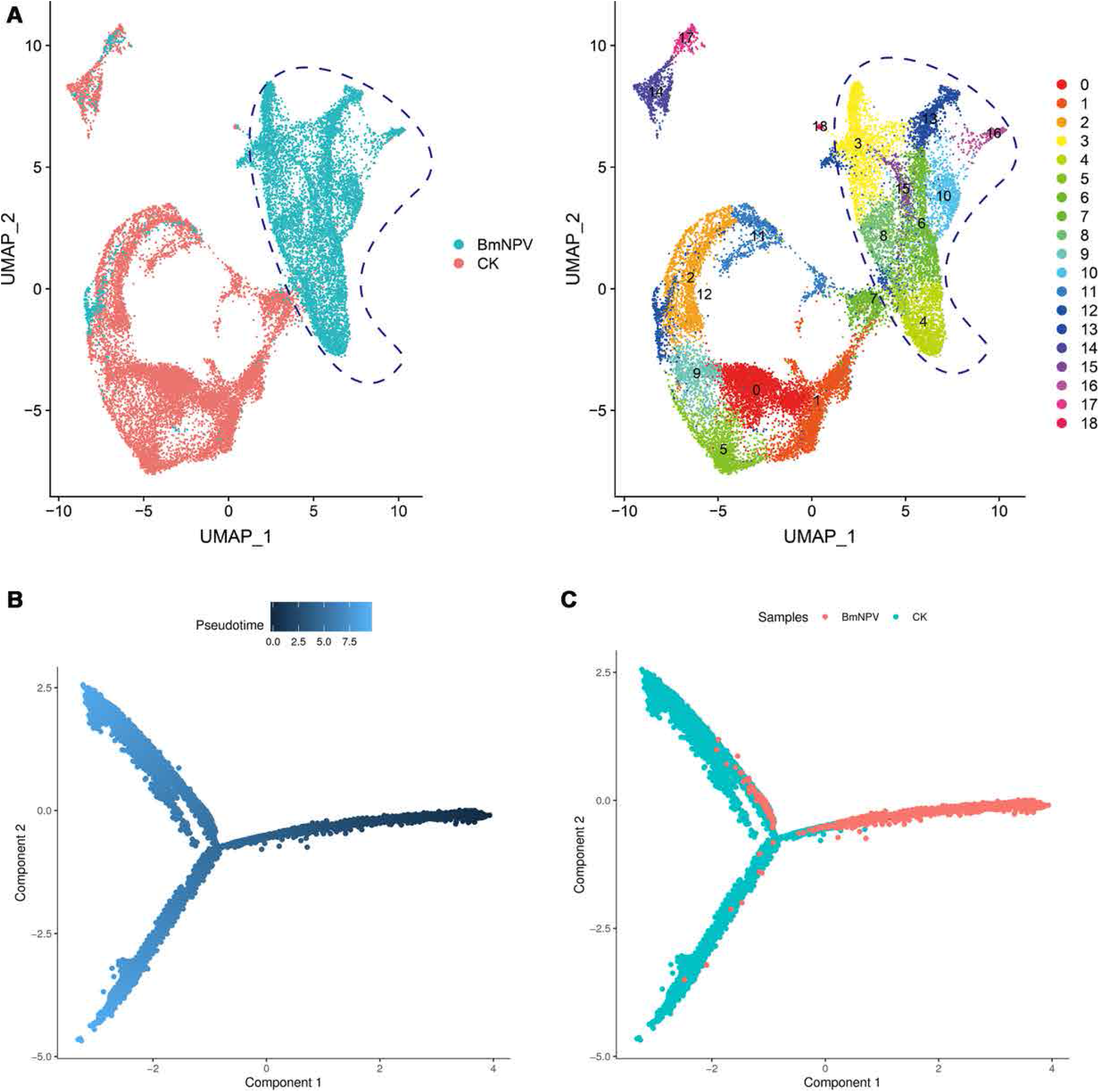
Hemocytes in the infection group are mainly located at the early stage of the pseudotime trajectories. **(A)** Uniform Manifold Approximation and Projection (UMAP) displaying all identified cells (left) and their subsets (right) in control and BmNPV-infected groups. **(B)** Pseudotime trajectory calculated from all cells in control BmNPV-infected groups. Darker dot colors represent shorter pseudotime and earlier differentiation period. **(C)** Mapping of hemocytes in control and BmNPV-infected groups to the pseudotime trajectory. Hemocytes in the infected group are mainly located at the early stage of the pseudotime trajectories.

### High viral load is a characteristic of major infected cell types

Since the viral mRNA is poly-adenylated, scRNA-seq can capture both viral and host mRNAs within each individual cell (Figure 1A). Individual BmNPV-infected cells could therefore be identified following alignment of viral genes with the BmNPV genome. Unexpectedly, viral genes can be detected in every cell of the BmNPV-infected sample (Supplementary file 5), illustrating widespread and efficient infection of hemocytes by baculovirus in the silkworm. In Figure 5A, the abundance of viral gene expression in individual cells is illustrated (values of the expression abundance of each viral gene are presented in Supplementary file 5). In most infected cells, high expression of viral genes is observed (Figure 5A). The top 20 of highly expressed BmNPV genes in each cell cluster are illustrated in Figure 5B and presented in detail in Supplementary file 5. The violin plots show that viral early genes (*ie-1* and *pe38*) as well as late genes (*vlf-1* and *lef-2*) and more specifically *vp39* (encoding the major capsid protein), *gp64/67* (envelope gene) and *gp41*(an essential gene required for the egress of nucleocapsids from the nucleus) (Rohrmann, 2019) are all highly expressed in most cells of each hemocyte cluster in the BmNPV-infected sample at 3 dpi (Figure 5C). When focusing on the distribution of the viral loads (defined as the proportion of viral UMIs in the total UMI content of a given cell) within infected cells of different hemocyte clusters, it was observed that infected cells in most of the clusters (0, 1, 3, 4, 6, 7, 8, 10 and 15) carried a high viral load of at least 20% (Figure 5D). Additionally, infected cells within cluster 2, 11, 12, 13, 14, 16, 17 and 18 displayed variability of viral-load, although the vast majority still maintained high viral-load states, except for those in cluster 14 (Figure 5D). Only in cluster 9, all infected cells carried a medium viral load (Figure 5D). Together, these data suggested that all hemocytes could be infected by BmNPV and that the majority of infected cells carried a high viral-load at 3 dpi.

**Figure 5.**
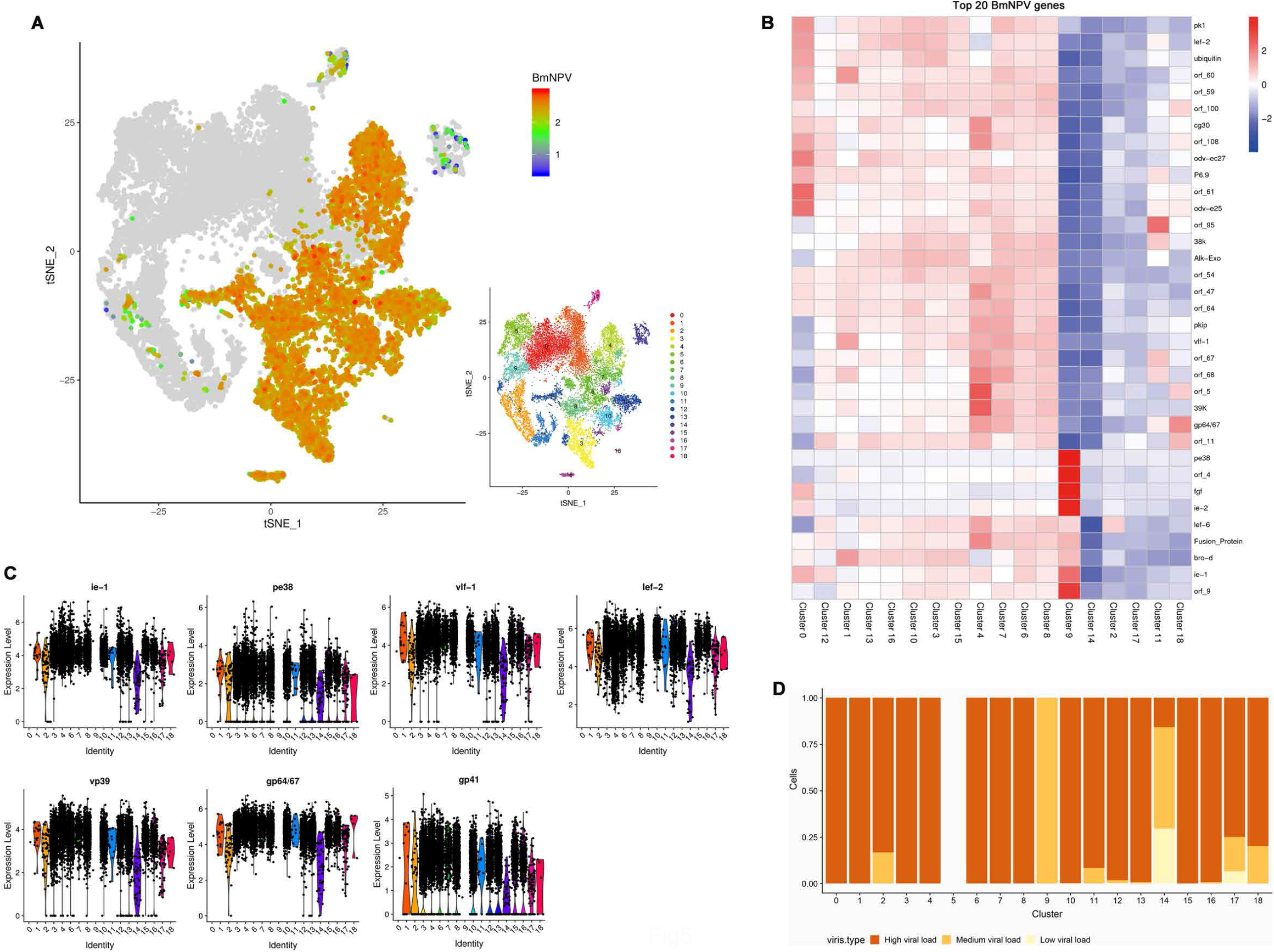
Analysis of viral gene expression and viral load in BmNPV-infected hemocytes. **(A)** t-SNE displaying the normalized expression (Z-score) of all viral genes in each BmNPV-infected cell. **(B)** Heatmaps showing the normalized expression (Z-score) of the top 20 of highly expressed BmNPV genes in each cell cluster of the BmNPV-infected group. **(C)** Expression levels of viral early genes (*ie-1* and *pe38*), late genes (*vlf-1* and *lef-2*) and characteristic viral genes (*vp39, gp64* and *gp41*) in hemocyte clusters. **(D)** The proportion of cells with different viral load in each cluster. Shown are the percentages of low (light yellow), medium (yellow), and high (brown) viral-load states (y axis) within the population of infected cells in each cluster (x axis).

### BmNPV infection suppresses the antiviral response in the major hemocyte cell types

To characterize the host responses against to BmNPV infection, we calculated the differential expression of genes between cell populations of the BmNPV-treated and control samples using Seurat (applied on clusters with more than 25 cells). Compared to the control sample, it was found that most of the host genes were down-regulated after BmNPV infection in all selected hemocyte clusters (2, 4, 6, 7, 14, 16 and 17) (Figure 6A; Supplementary file 6; Figure supplement 3). This result suggests that the defense ability in most of BmNPV-infected cells was already compromised at the later stages of infection. When GO and KEGG enrichment analysis of DEGs was performed for each cluster, the GO terms “response to stimulus” and “immune system process” and the KEGG pathways “Immune system”, “Infectious diseases” and “Signal transduction” were enriched with the down-regulated DEGs (Figure supplement 4). On the other hand, 243 DEGs were induced after BmNPV infection in cluster 2 (Figure 6A; Supplementary file 6; Figure supplement 3), indicating that this cluster may play an important role in the host antiviral response at the later stage of infection. However, the cells in cluster 2 were significantly depleted after BmNPV infection (Figure 3B, D), therefore questioning the relevance of their contribution to overall host protection. The picture prevails that in the late stage of BmNPV infection the virus completely overcomes the host defenses.

**Figure 6.**
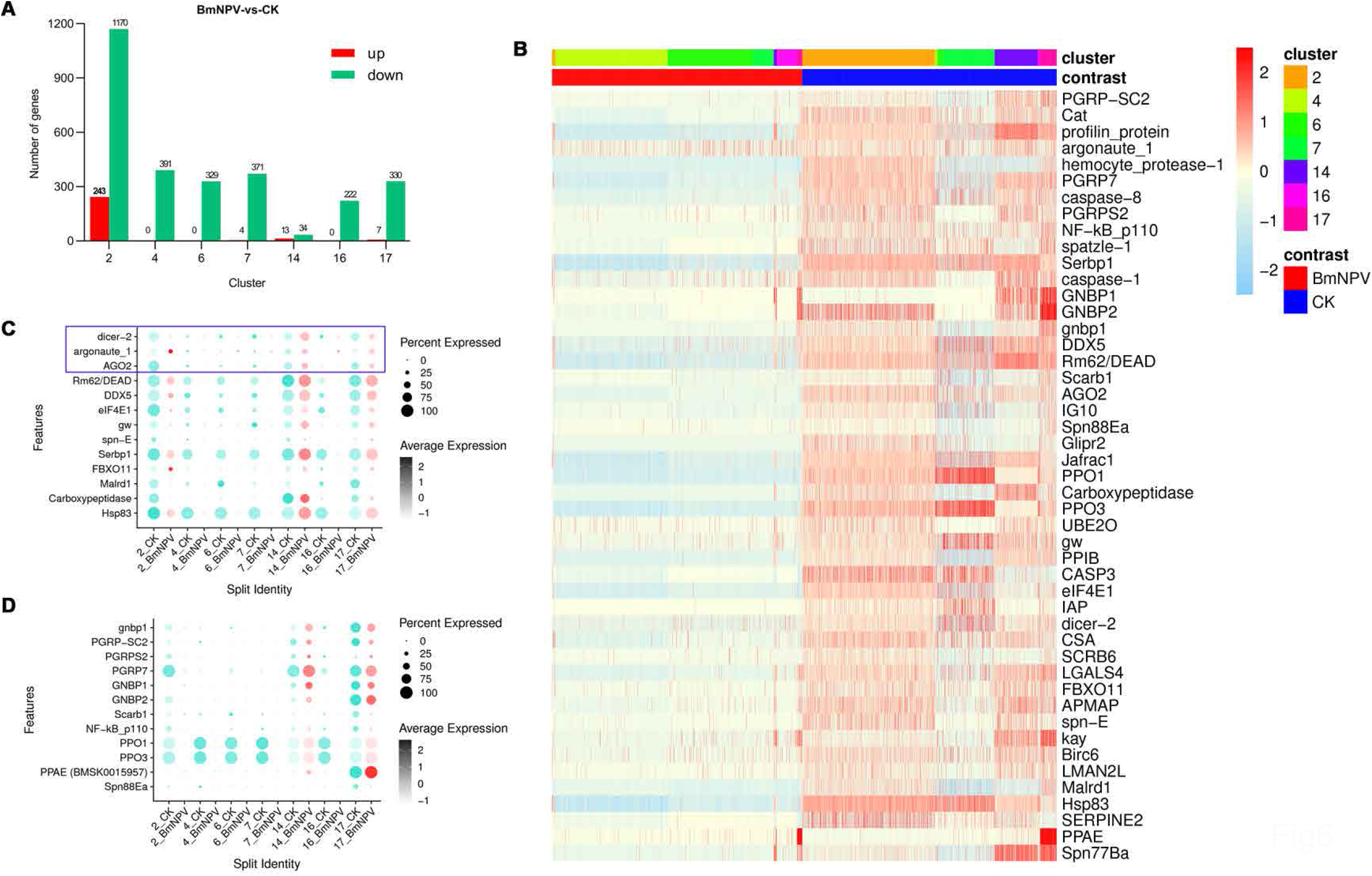
DEGs analysis between BmNPV-infected and control cells within cell clusters. Seurat was used for DEGs analysis for clusters with more than 25 cells. **(A)** Histogram showing all of up-regulated (red) and down-regulated DEGs (green) in BmNPV-infected cells compared to control cells within selected clusters. **(B)** Heatmap showing the normalized expression (Z-score) of the immune-related DEGs in various cells of hemocyte clusters: cells from the BmNPV-infected sample are compared with the control sample. **(C, D)** Dot plot representing DEGs belonging to “RNAi” (C) and “innate immune pathway” (D) categories in hemocyte clusters: cells from the BmNPV-infected sample are compared with the control sample. The black gradient represents the expression level, while the size of the dots represents the percentage of cells expressing any gene per cluster. Red dots represent the BmNPV-infected group and green dots represent controls.

The landscape of immune-related DEGs shows mainly inhibition in hemocyte clusters 2, 4, 6, 7, 14, 16 and 17 during BmNPV infection (Figure 6B). Specifically, DE RNAi-related genes are mainly down-regulated in the BmNPV-infected sample (Figure 6C). It is worth noting that the genes encoding Dicer-2 and Argonaute-2, the core factors of the siRNA pathway, did not change in expression to BmNPV infection at 3 dpi (Figure 6C). Additionally, the DEGs involved in pathogen pattern recognition and innate immune pathways are also inhibited in the BmNPV-infected sample (Figure 6D). Thus, we speculate that on the third day after infection with BmNPV, the host’s antiviral system in hemocytes is almost in a state of collapse.

### Identification of specific cell types in the hemolymph of silkworm

Since there are no recognized marker genes for various types of silkworm hemocytes, it was not obvious to classify the identified clusters 0-18 into the specific subgroups described in the literature such as plasmatocytes (PL), granulocytes (GR), spherulocytes (SP) and oenocytoids (OE) (Lavine and Strand, 2002). However, we tried to distinguish clusters 0-18 based on the genes that are highly expressed in each subgroup of silkworm hemocytes determined by morphology in the literature. These potential marker genes found in the literature (Nakahara et al., 2009, Zhang et al., 2014, Zhang et al., 2018) were subjected to the BLAST function of SilkDB 3.0 (Lu et al., 2020) to obtain the new gene IDs of the database. Specifically, Nakahara *et al*. (Nakahara et al., 2009) determined that *serine protease homolog 1*(BMSK0013968), *βGRP3* (BMSK0006299) and *paralytic peptide* (BMSK0007609) were expressed only in plasmatocytes and no other hemocyte subset; that t*SCR-C* (BMSK0015652), *CatB* (BMSK0010120), *HP1* (BMSK0003908) and *PGRP* (BMSK0004739) were detected in granulocytes; that *PPBP1* (BMSK0014159) and *PPBP2* (BMSK0013765) were strongly expressed in oenocytoids; and that *cathepsin L-like protein* (BMSK0005696) can be used as a spherulocyte-specific molecular marker in silkworm. Moreover, Kui Zhang *et al* reported that *integrinaPS3* (BMSK0006871) could be used as a specific marker for silkworm granulocytes (Tattikota et al., 2020). Additionally, in Kui Zhang’s doctoral thesis (In Chinese, 2017), they found that *Integrin β3* (BMSK0001792) and *BmSCRB8* (BMSK0013677) were strongly expressed in plasmatocytes and granulocytes, respectively. Moreover, the promoter of *Integrin β3* can drive EGFP specifically in plasmatocytes but not in other hemocyte types (Zhang et al., 2018).

We explored the expression of the genes mentioned in literature in clusters 0-18 and displayed them in bubble plot (Figure 7A). Based on these potential marker genes, we first identified cluster 18 as spherulocytes (SP) based on *cathepsin L-like protein* (BMSK0005696) expression (Figure 7A, B). Next, we assigned cluster 14 and 17 as plasmatocytes because of the presence of DGEs *serine protease homolog 1*(BMSK0013968), *paralytic peptide* (BMSK0007609), *βGRP3* (BMSK0006299) and *Integrin β3* (BMSK0001792) (Figure 7A, B). We further assigned cluster 0, 1,2,5 and 9 as granulocytes because the presence of markers such as *SCR-C* (BMSK0015652), *CatB* (BMSK0010120), *HP1*(BMSK0003908), *PGRP* (BMSK0004739), *BmSCRB8* (BMSK0013677) and *integrinaPS3* (BMSK0006871) (Figure 7A, B). Finally, clusters 7 and 11 were identified as oenocytoids because of high expression of *PPBP1* (BMSK0014159) and *PPBP2* (BMSK0013765) (Figure 7A, B). Moreover, the Pearson’s Correlation analysis between bulk RNA-seq of partially purified granulocytes and plasmatocytes and scRNA-seq showed that clusters 0, 1, 2, 5 and 9 have high correlation with granulocytes and that clusters 14 and 17 likely correspond with plasmatocytes (Figure 7C). This result increases the reliability of the assignment of scRNAseq clusters as plasmatocytes and granulocytes in this study.

**Figure 7.**
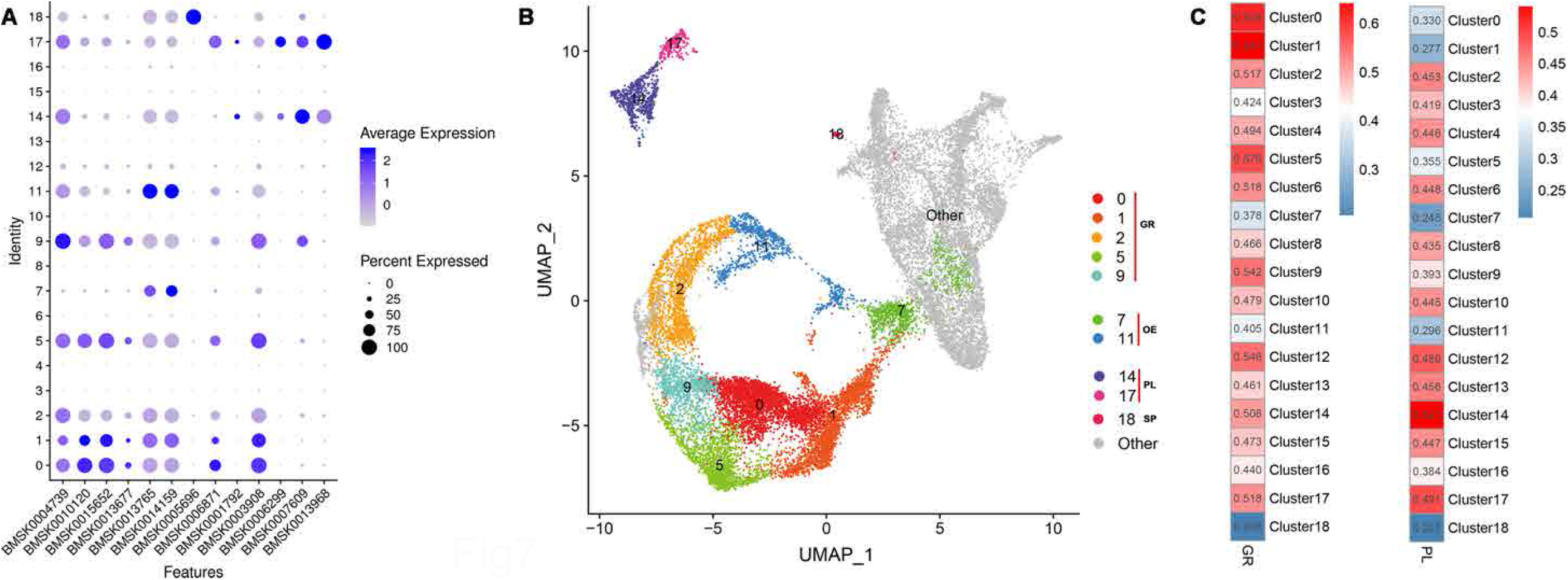
Specific cell types in the hemolymph of silkworm larvae. **(A)** Potential marker genes collected from published literature and their expression in hemocyte clusters 0 to 18. Gene identities of SilkDB3.0 correspond to the following genes in literature: BMSK0004739 (*PGRP*), BMSK0010120 (*CatB*), BMSK0015652 (*SCR-C*), BMSK0013677 (*BmSCRB8*), BMSK0013765 (*PPBP2*), BMSK0014159 (*PPBP1*), BMSK0005696 (*cathepsin L-like protein*), BMSK0006871 (*integrinaPS3*), BMSK0001792 (*Integrin β3*), BMSK0003908 (*HP1*), BMSK0006299 (*βGRP3*), BMSK0007609 (*paralytic peptide*), BMSK0013968 (*serine protease homolog 1*). The colour gradient of the dot represents the expression level and the size reflects the percentage of cells expressing these genes per cluster. **(B)** UMAP displaying the plasmatocytes (PL), granulocytes (GR), spherulocytes (SP), oenocytoids (OE) and unclassified cells (Other – mainly in BmNPV-infected group). **(C)** Pearson’s Correlation analysis between bulk-RNAseq data from partially purified GR and PL and scRNA-seq data of different cell clusters based on gene expression levels.

Granulocytes, plasmatocytes, spherulocytes, and oenocytoids are considered as the four main circulating haemocyte types in Lepidopteran species (Ribeiro and Brehelin, 2006). In particular, granulocytes and plasmatocytes together usually comprise more than 50% of the hemocytes in circulation at the larval stage (Lavine and Strand, 2002). In our analysis, an effort was made to assign the different hemocyte clusters in scRNAseq to the four major cell types (Figure 7). However, these four main hemocyte types become strikingly depleted in the hemolymph of BmNPV-infected silkworms (Figure 3B, D; Figure 4A; Figure 7B) and are replaced by groups of cells at the early stage of differentiation in the pseudo-timeline (Figure 4). Prohemocytes are hypothesized to be progenitors that differentiate into one or more of the other hemocyte types (Nakahara et al., 2010, Yamashita and Iwabuchi, 2001). It can therefore be speculated that hemocytes of clusters 3, 4, 6, 8, 10, 12, 13, 15 and 16, that appear mainly in the BmNPV-infected silkworm, could correspond to progenitor cells that are released from the hematopoietic organs (Liu et al., 2013) as a response to hemocyte depletion during BmNPV infection.

## Discussion

Hemocytes, the motile cells that populate the hemolymph in insects, can carry out multiple and diverse functions. Similar to macrophages in vertebrates, differentiated hemocytes play an important role in body homeostasis through tissue repair (e.g. production of extracellular matrix around differentiating tissues) and disposal of cellular waste (e.g. phagocytosis of debris from apoptotic cells) (Hartenstein, 2020). In addition, hemocytes play a crucial role in both cellular (e.g. phagocytosis of microbial pathogens, encapsulation of large structures such as eggs from parasitoid eggs) and humoral immunity (e.g. production of AMPs in response to activation of immune signaling pathways) (Browne et al., 2013). Hemocytes are also involved in the uptake and storage of lipids and nutrients (Cattenoz et al., 2020).

To understand their diverse functions, a detailed classification of the different types of hemocytes is necessary. With respect to Lepidoptera, morphological and biochemical properties have led to the designation of five major hemocyte cell types (mentioned above) (Lavine and Strand, 2002, Liu et al., 2013) which should be considered only as a very rough approximation for the expected much higher diversity at the molecular level. In *Drosophila*, only three major hemocyte types are distinguished (plasmatocytes, crystal cells, lamellocytes) that were correlated to the five types in Lepidoptera (Liu et al., 2013, Lavine and Strand, 2002). However, compared to Lepidoptera, genetic characterization of hemocyte lineages in *Drosophila* is much more advanced and clear molecular markers for the three different lineages were identified (Buchon et al., 2014, Honti et al., 2014). ScRNA-seq analysis of hemocytes confirmed the three lineages in *Drosophila*, although a much higher complexity was revealed at the molecular level. More specifically, the broad categories of plasmatocytes, crystal cells and lamellocytes were found to be distributed into 12, 2 and 2 cell clusters, respectively, that are thought to reflect functional diversification within a broad cell type or represent different stages of maturation (Tattikota et al., 2020, Cattenoz et al., 2020). Attempts were made to classify hemocytes in Lepidoptera at the molecular level, e.g. through the use of specific antibodies, but their molecular targets were never elucidated (Beetz et al., 2004, Gardiner et al., 1999).

In this work, the technique of scRNA-seq was applied to categorize different hemocyte populations in 5^th^ instar larvae of the silkworm that were either mock-infected (injection with PBS) or infected with a high dose of BmNPV for a period of three days. Nineteen different hemocyte clusters were identified during joint analysis of the scRNA-seq data from both treatments revealing high heterogeneity among the hemocytes that populate the hemolymph in larvae (of note, when scRNA-seq of the two samples were analyzed separately, highly similar results were obtained, reflecting the robustness of the approach that was used – see Supplementary Information). Strikingly, cell clusters that are prevalent in the BmNPV-infected group (3, 4, 6, 8, 10, 13, 15 and 16) were clearly separated from clusters that were specific for the control group (0, 1, 2, 5, 9, 11 and 18). In only two clusters (7 and 17), the amount of cells was divided roughly equally between the infected and uninfected condition. Thus, baculovirus infection clearly has a major impact on the hemocyte population after three days of infection.

Our analysis and division of hemocytes in different clusters allowed to assign particular “marker genes” (top 5 of differentially expressed genes) to each cluster (Figure 1). However, the number of differentially expressed genes was much higher in uninfected cells than in infected cells, revealing the impact of baculovirus infection. For the uninfected condition, the “marker genes” in the different clusters could give important clues about the functional specialization of the cells that belong to the cluster. Based on the analysis of data in the literature, we have tentatively assigned the different clusters to the particular cell types that were described previously during morphological and cytochemical observations (i.e. prohemocytes, plasmatocytes, granulocytes, oenocytoids, spherulocytes) (Figure 7).

On the other hand, it is rather difficult to assign a particular function to a specific cell cluster based on the top 5 of differentially expressed “marker genes” that emerged from the analysis. To give an example, the highest ranked genes in cluster 9 encode (1) Imaginal Morphogenesis Protein-Late 2 (IMP-L2), a secreted factor and member of the immunoglobulin family (Garbe et al., 1993); (2) the antimicrobial peptide Cecropin B (Kato et al., 1993); (3) mannosyl-oligosaccharide alpha-1,2-mannosidase IA that participates in the maturation process of N-glycans during protein glycosylation in the Golgi complex (Jarvis et al., 1997); (4) unknown protein LOC106126609 (HHpred (Zimmermann et al., 2018) shows a small region with homology to ubiquitin-binding proteins) and (5) another unknown protein of which the sequence is likely incomplete but with a region of resemblance to membrane channels that selectively transport water and small neutral and ionic molecules (e.g. aquaporins). IMP-L2 binds insulin-like peptides and functions as an antagonist of insulin/insulin-like growth factor signaling (Sarraf-Zadeh et al., 2013). As such, IMP-L2 regulates developmental timing and could be involved in the regulation of aerobic glycolysis in hemocytes during the immune response (Krejcova et al., 2019).

The high expression of *Cecropin B* genes (6 genes belonging to the same cluster on chromosome 26 with log2FC values of 2.9, 2.8, 2.6, 2.4, 2.2, and 2.0) in the uninfected condition may reflect a regulatory function in developmental processes such as cuticle formation via the regulation of prophenoloxidase expression (Liu et al., 2017) (as opposed to a defense against bacterial infection). The increased expression of IMP-L2 and Cecropins indicates that cluster 9 may be associated with humoral immunity or have a regulatory function by the secretion of cytokines. To support the latter, it is noted that paralytic peptide, a cytokine with diverse functions in growth and immunity (Matsumoto et al., 2012), is also upregulated in cluster 9 (log2FC = 2.6). Also the increased expression of a gene associated with protein glycosylation, which is most often associated with secreted and membrane proteins, is conform to such function. With respect to the potential expression of an aquaporin genes, it is noted that in *Drosophila*, the aquaporin Prip is one of the most enriched genes in plasmatocytes (Ramond et al., 2020) and another aquaporin, Drip, is enriched in lamellocytes (Tattikota et al., 2020). During cellular immunity in *Spodoptera exigua*, aquaporins are involved in the regulation of cell shape (Ahmed and Kim, 2019).

The above analysis indicates that the identification of particular marker genes only gives a general indication of the possible function of the cells in a particular cluster, especially if it involves an intermediary cell type (as seems to be the case for cluster 9). To solve this conundrum, future studies should try to correlate the morphology of individual cells with their transcriptomes, for which, however, technological advances are needed. Another approach could be to monitor the changes in abundance of cell clusters during development or during infection of different pathogens which could reveal their importance during different physiological and pathological states.

It can be noted that for a few clusters their function is clearer. For instance, all the identified marker genes in cluster 5 (regulator of microtubule dynamics, kinesin-like protein, spindle assembly checkpoint proteins) are related to proliferation and mitosis, indicating their correspondence to prohemocytes or proliferating plasmatocytes. Clusters 17 and 18, which are the least abundant, clearly represent terminally differentiated cells because of the very high induction of a subset of genes (9 and 16 genes with log2FC > 8, respectively). Interestingly, the gene with the highest differential expression in cluster 18 encodes one of the early class “CA” chorion proteins, which were thought to be exclusively expressed in the ovary and are constituents of the eggshell (Chen et al., 2015). Early chorion proteins are proposed to form an initial scaffold during the first steps of chorion assembly and their expression in cluster 18 hemocytes may reflect a similar role in the assembly of another protective layer, the cuticle. In this respect, it should be remembered that the control group in this study was treated with PBS through injection and that the small group of cluster 18 (47 cells) could have been involved in wound repair.

Baculovirus infections are well known for their virulent character and their large impact on cellular physiology (Nguyen et al., 2013b). During baculovirus infection of tissue culture cells, host mRNAs decrease to less than 10% after 48 hr while total (mainly viral) mRNAs increase by 70% (Nguyen et al., 2013a). In the baculovirus expression vector system, host genes that show an increase in expression relate to the stress response to unfolded proteins indicating that the cellular expression capacity is at its limit (Koczka et al., 2018). However, most studies that monitor the transcriptional response of the host cells to baculovirus infection investigate mainly early responses and are limited to 48 hr post-infection (Nguyen et al., 2013b), while in this study, scRNA-seq was applied on hemocyte samples after 3 days of infection. Cell clusters that are dominated by infected cells seem to fall in two groups: clusters 4, 6, 8, 10 and 12 have none to few DGEs with low log2FC values, while clusters 3, 13, 15 and 16 show high induction of a limited number of genes that are cluster-specific such as unknown genes (cluster 3), actin cytoskeleton regulatory protein and serine proteases (cluster 13), neuropeptide-like precursor 4E (cluster 15) and motility genes (myosin, troponin, actin; cluster 16). Clusters 3, 6, 8, 10, 13, 15 and 16 all seem related because of overlapping DGEs (cuticular protein RR-2 motif, Ser/Arg-rich splicing factor, serine protease precursor, trypsin-like protease and others). On the other hand, DGEs in cluster 12 are totally different (while having very low log2FC) and no DGEs were identified in cluster 4.

Pseudo-time analysis reveals that the BmNPV-infected cells are at the early stages of the calculated trajectories (Figure 4) which could lead to the hypothesis that they correspond to stem cells or prohemocytes. However, this assumption seems to contradict with the analysis of the DGEs in the cell clusters that are prevalent in the BmNPV-infected sample. DGEs in these cell clusters (as discussed above) generally are representative of specialized functions which is not expected from progenitor cells. Baculoviruses routinely transform infected cells into “virus production factories” and shut-off of host gene expression can be assumed to cause a loss of differentiation features (including a decrease in expression of immune-related genes). Loss of host gene expression in infected cells could have caused their artefactual assignment as “early stage” during pseudo-time modeling calculations. Taking into account the above caveat, it is nevertheless considered that the limited number of DEGs could be a hallmark of “resting” precursor cells that are prevented from maturation by baculovirus infection.

Based on the results of this research, we present the following hypothetical timeline of the natural infection process of silkworms by BmNPV: first, BmNPV occlusion bodies are ingested by the silkworm and become dissolved in the midgut, and occlusion derived virions (ODV) are released which then infect midgut epithelial cells (Figure 8A). Subsequently, budded viruses (BV) are produced which spread the infection to the entire silkworm including all hemocytes (Figure 8B). Viral infection causes severe depletion of the main silkworm hemocyte subgroups including Granulocytes (GR), Plasmatocytes (PL), Oenocytoids (OE) and Spherulocytes (SP) (Figure 8C). Furthermore, the host immune responses in silkworm hemocytes were inhibited by BmNPV infection at the late stage (Figure 8D). In order to replenish the lost hemocytes including GR, PL, OE and SP, the host mobilizes a large number of progenitor cells such as prohemocytes into the circulating hemolymph. However, after the virus infects these progenitor cells, it may prevent them from continuing to differentiate into other types of hemocytes. Therefore, the exhaustion of all blood cells is inevitable (Figure 8E). Due to the exhaustion of the main hemocyte subsets and the severe suppression of the immune response, the silkworm’s antiviral system collapses causing death (Figure 8F).

**Figure 8.**
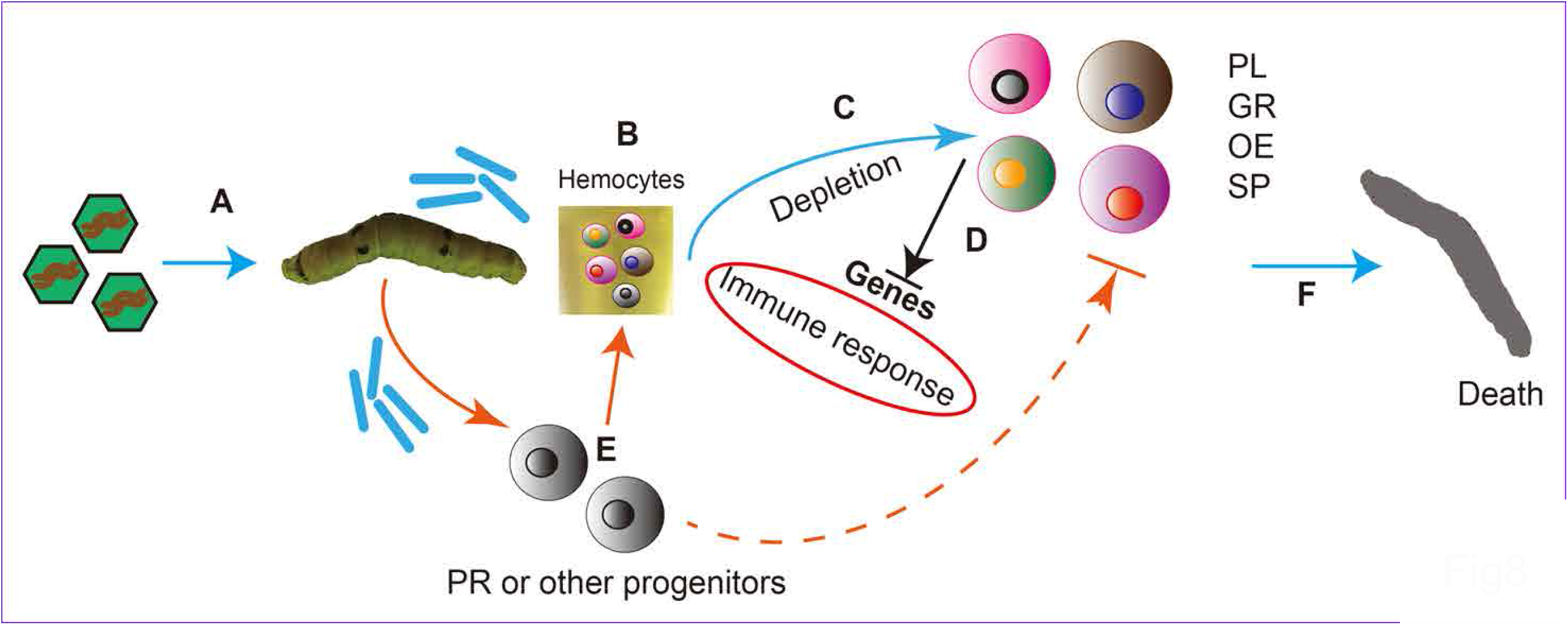
Hypothetical timeline of the natural infection process of silkworms by BmNPV. **(A)** BmNPV occlusion bodies are ingested by the silkworm and become dissolved in the midgut, and occlusion derived virions (ODV) are released which then infect midgut epithelial cells. **(B)** Budded viruses (BV) are produced which spread the infection to the entire silkworm including all hemocytes. **(C)** Viral infection causes severe depletion of the main silkworm hemocyte subgroups including Granulocytes (GR), Plasmatocytes (PL), Oenocytoids (OE) and Spherulocytes (SP). **(D)** The host immune responses in silkworm hemocytes were inhibited by BmNPV infection at the late stage. **(E)** In order to replenish the lost hemocytes including GR, PL, OE and SP, the host mobilizes a large number of progenitor cells such as prohemocytes into the circulating hemolymph. However, after the virus infects these progenitor cells, it may prevent them from continuing to differentiate into other types of hemocytes. Therefore, the exhaustion of all blood cells is inevitable. **(F)** Due to the exhaustion of the main hemocyte subsets and the severe suppression of the immune response, the silkworm’s antiviral system collapses causing death.

In summary, our scRNA-seq study on hemocytes in the silkworm represents a rich resource of data that can be mined for future experiments to address the function of these cells during development and in response to pathogenic infection in Lepidoptera. The “marker genes” that were assigned to different clusters need to be confirmed by independent methods such as *in situ* hybridization and immunocytochemistry with specific antibodies. With respect to the control group, experimental manipulation (injection and wounding) and developmental stage (following the molt) could have made an impact on the composition of hemocytes in the analysis of the scRNAseq data (i.e. biased towards tissue remodeling). Checking hemocyte populations at different developmental stages or after particular experimental interventions (e.g. RNAi, hormones, cytokines) will lead to additional insights. Regarding BmNPV infection, earlier time points or lower doses of virus may provide new insights into the vulnerability of particular hemocyte types and the mounting of an immune response. Our study confirms that the technique of scRNA-seq provides a great advancement for the study of hemocytes as well as other tissue types and will lead to a more complete understanding of biological processes and organ function.

## Materials and Methods

### Silkworm and BmNPV infection

Larvae of silkworm (*B. mori*, Dazao strain) were reared with fresh mulberry (*Morus*) leaves and reared under constant environmental conditions of 28 °C and 60-70% humidity. Recombinant BmNPV-EGFP (Enhanced Green Fluorescent Protein), as a reporter virus, was constructed by Bac-to-Bac Baculovirus Expression System and kept in Guangdong Provincial Key Laboratory of Agro-animal Genomics and Molecular Breeding. Newly molted fifth-instar silkworm larvae were injected with either 10μL of BmNPV-EGFP (10^5.8^ TCID_50_/0.1 mL) or phosphate-buffered saline (PBS) (injection control). Blue Light Gel Imager (Sangon Biotech, China) was used to monitor the spread of green fluorescence in infected larvae on a daily basis until whole body expression was achieved.

### Preparation of single hemocytes in suspension

At three days after BmNPV infection, three mL of hemolymph was collected from pools of twenty BmNPV-infected animals and twenty controls on ice and centrifuged at 500 g for 2 min at 4 °C. The hemocyte pellet was washed twice using 1 mL cold Grace’s insect medium (Gibco, USA) supplemented with 10% FBS (Gibco, USA), which was followed by filtering through a 40 µm cell strainer (Biosharp, China) and centrifugation at 500 g for 2 min at 4 °C. The hemocyte pellet was resuspended using 200 µL PBS (calcium and magnesium-free, Gibco, USA) supplemented with 0.04% bovine serum albumin (BSA) (Solarbio, China). Cell viability and number were assessed by 0.4% trypan blue staining and cell counting using a hemocytometer. High quality single hemocyte preparations were subjected to encapsulation by 10X Genomics v3 kit (10x Genomics, USA).

### Single hemocyte encapsulation and sequencing

Single cell encapsulation, complementary DNA (cDNA) library synthesis, RNA-sequencing and data analysis were completed by Gene Denovo (Guangzhou, China). The single-cell suspensions were bar-coded and reverse-transcribed into scRNA-seq libraries using the Chromium Single Cell 3’ Gel Bead-in Emulsion (GEM), Library and Gel Bead Kit (10x Genomics, USA) according to the manufacturer’s protocol. Briefly, single silkworm hemocytes were barcode-labeled and mixed with reverse transcriptase into GEMs, then the cDNA library was amplified using PCR with the sequencing primers R1 and R2, and subsequently ligated to Illumina sequencing adapters with P5 and P7. Finally, the cDNA libraries were sequenced on the Illumina 10x Genomics Chromium platform (Illumina Novaseq 6000).

### ScRNA-seq data processing and analysis

The latest published version of the silkworm genome (SilkDB3.0) was used in the present study (Lu et al., 2020). Since SilkDB3.0 does not contain mitochondrial genome, we have added silkworm mitochondrial genome data (NCBI Reference Sequence: NC_002355.1) in the subsequent analysis. The raw scRNA-seq data were aligned, filtered, and normalized using Cell Ranger (10x Genomics) software (Cell Ranger 3.1.0). A read is considered as exonic if at least 50% of it intersects an exon, intronic if it is non-exonic and intersects an intron, and intergenic otherwise. A read that is compatible with the exons of an annotated transcript, and aligned to the same strand, is considered mapped to the transcriptome. If the read is compatible with a single gene annotation, it is considered uniquely (confidently) mapped to the transcriptome. Only reads that are confidently mapped to the transcriptome are used for Unique Molecular Identifier (UMI) counting.

Seurat is a popular R package that can perform quality control (QC), analysis, and exploration of scRNA-seq data, which was originally developed as a clustering tool for scRNA-seq data (Satija et al., 2015). In this study, the cell subset was grouped by graph-based clustering based on the gene expression profile of each cell in Seurat. The data were clustered using principal component analysis (PCA) and visualized with t-distributed Stochastic Neighbor Embedding (t-SNE) (Linderman et al., 2019) or Uniform Manifold Approximation and Projection (UMAP) (Becht et al., 2019). Other data analyses including standardization, difference of gene expression, and marker gene screening were also achieved by Seurat.

Notably, scRNAseq data were validated for analysis according to the following QC criteria: 1) 200 < gene counts < 4000 per cell; 2) UMI counts < 30, 000 per cell; 3) percentage of mitochondrial genes < 25%. After normalizing the data using Seurat, gene expression level in scRNA-seq was calculated as log(1 + (UMI A ÷ UMI Total) × 10000).

### Differentially expressed genes (DEGs) analysis per cluster

Likelihood-ratio test (McDavid et al., 2013), performed on single cluster cells against all other cells, was used to identify differentially expressed genes (DEGs) in single clusters based on differential expression. Up-regulated DEGs in each cluster were identified the following criteria: 1) p value ≤ 0.01; 2) log2FC ≥ 0.360674 (log2FC means log fold change of the average expression between the two groups); and 3) percentage of cells in a specific cluster where the gene is detected > 25%. Gene Ontology (GO) and Kyoto Encyclopedia of Genes and Genomes (KEGG) pathway enrichment analysis were further carried out based on gene expression levels to identify the main features of each cluster. The top 5 genes in each cluster were selected as the potential marker genes according to the result of DEGs analysis. Then the expression distribution of each marker gene among clusters was visualized by bubble diagrams.

### Pseudo temporal ordering of cells

Single cell trajectory was analyzed using a matrix of cells and gene expression levels by Monocle 2 (Version2.6.4) (Trapnell et al., 2014). Monocle reduces the space in which the cells were embedded to two dimensions and orders the cells (parameters used: sigma= 0.001, lambda = NULL, param.gamma = 10, tol= 0.001). Once the cells were ordered, the trajectory (with tree-like structure, including tips and branches) could be visualized in the reduced dimensional space.

### Viral genes analysis in cell clusters of the BmNPV-infected group

To determine the state of BmNPV infection in each hemocyte cluster, the expression of viral genes and viral load were analyzed in the different cell clusters in the BmNPV-infected group. Viral gene expression was analyzed using Cell Ranger, based on the BmNPV genome (GenBank: JQ991008.1). The ‘viral load’ of a cell in scRNA-seq analysis was based on the number of UMIs that map to the BmNPV genome and expressed as a percentage of total UMI content of a given cell (Steuerman et al., 2018). In the present study, infected hemocytes were divided into three categories of viral-load states: low (< 5%), medium (between 5% and 20%), and high (> 20%).

### DEGs analysis in cell clusters between BmNPV-infected and uninfected groups

To explore the response of each hemocyte cluster to BmNPV, we further analyzed the DEGs between the BmNPV-infected group and the control group using Seurat software under the condition of a minimum of 25 cells per cluster. New identities were set up as “group_cluster” or “group_cell type” for analysis (Stuart et al., 2019). A hurdle model in MAST (Model-based Analysis of Single-cell Transcriptomics) (Finak et al., 2015) was used to find DEGs for a group in one cluster. DEGs between the BmNPV-infected and control groups were identified by the following criteria: 1) |log2FC| ≥0.25; 2) p value_adj ≤ 0.05; and 3) percentage of cells where the gene is detected in a specific cluster > 25%. Identified DEGs were subsequently subjected to GO and KEGG pathway enrichment analysis.

### Analysis of bulk RNA-seq data

Bulk RNA-seq raw data of silkworm plasmatocytes (PL) and granulocytes (GR) were kindly provided by Dr Kui Zhang and Prof Hongjuan Cui (unpublished data, Southwest University, China). Plasmatocytes used in their study were obtained by culturing hematopoietic organs (from *B. mori*, Dazao strain) *in vitro* and granulocytes (GR) were obtained by culturing the adherent cells collected from the Dazao silkworm from which the hematopoietic organs were removed (Kui Zhang’s doctoral thesis, in Chinese, 2017). Since the flow cytometry sorting method cannot be used to separate silkworm hemocytes due to the lack of known marker genes and corresponding interacting specific antibodies, the PL and GR isolates described above are therefore considered to be impure preparations.

After removing low-quality reads and ribosomal RNA, high quality clean reads were mapped to the latest version of the silkworm genome (SilkDB 3.0) using HISAT2. 2.4 (Kim et al., 2015). The mapped reads were assembled using StringTie v1.3.1 (Pertea et al., 2016). FPKM (fragment per kilobase of transcript per million mapped reads) values were calculated to quantify genes’ expression abundance and variations, using StringTie software (Pertea et al., 2016). Pearson’s correlation analysis was used to investigate the relationships between bulk RNA-seq data of PL, GR and hemocyte clusters in scRNA-seq based on levels of gene expression.

## Acknowledgments

Thanks to Prof Hongjuan Cui and Dr. Kui Zhang (Southwest University) for kindly providing the raw data of the bulk RNA-seq. Thanks are due to Gene Denovo Corp. for its help in bioinformatics analysis. We also thank Jingjing Ning for her help in project coordination.

## Author contributions

MF and LS participated in the design of the study, collected and analyzed data and drafted the manuscript. JX, SF, XW, YZ and PW helped with sample preparation and data analysis. JS participated in the design and coordination of the study, and revised the manuscript. All authors read and approved the final manuscript.

## Data availability

Sequencing data have been deposited in BioProject under the accession number PRJNA658439_∘_

---

**Appendix File 1. scRNA-seq data statistics**.

**Appendix File 2. Up-regulated differentially expressed genes (DEGs) in each silkworm hemocyte cluster**.

**Appendix File 3. Top 5 DEGs of each silkworm hemocyte cluster**.

**Appendix File 4. Immune-related DEGs selected from clusters 0 to 18 and their distribution in each cluster**.

**Appendix File 5. Statistics of virus-infected cells and expression of viral genes in the different clusters of the BmNPV-infected group**.

**Appendix File 6. Properties of identified DEGs in clusters 2, 4, 6, 7, 14, 16 and 17 between the BmNPV-infected and control groups**.

**Appendix Fig S1.**
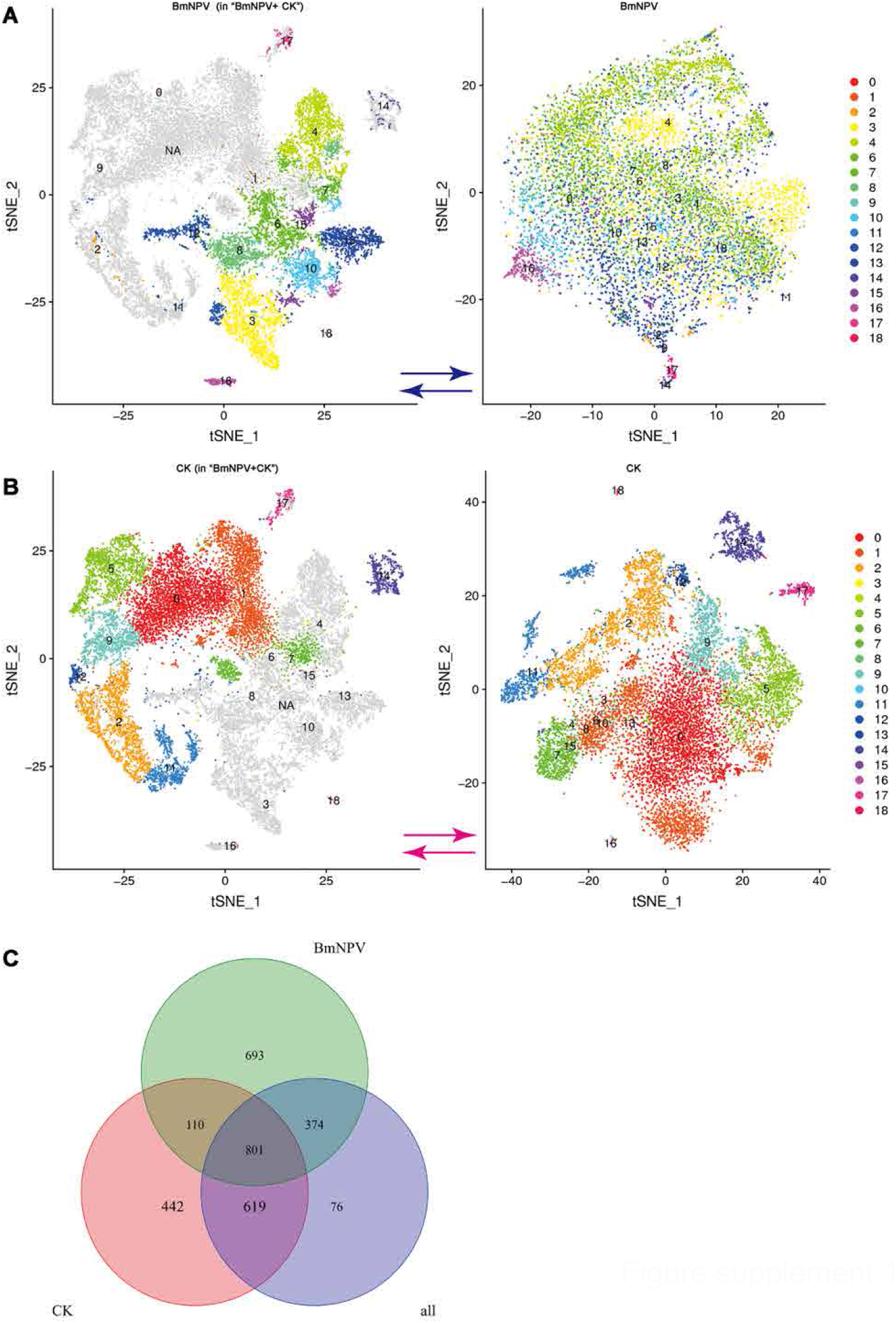
The analysis of hemocytes of the separate groups of control and BmNPV infection. **(A, B)** The hemocytes of the separate groups of BmNPV infection (A) and control (B) were mapped to the unified set (BmNPV+CK) using cell-specific barcodes. The agreement between the two clustering approaches is relatively high, especially for the cells in the control group (B). (C) The hypervariable genes that were identified for cell cluster classification in scRNA-seq were compared between individually analyzed groups and the unified set.

**Appendix Fig S2.**
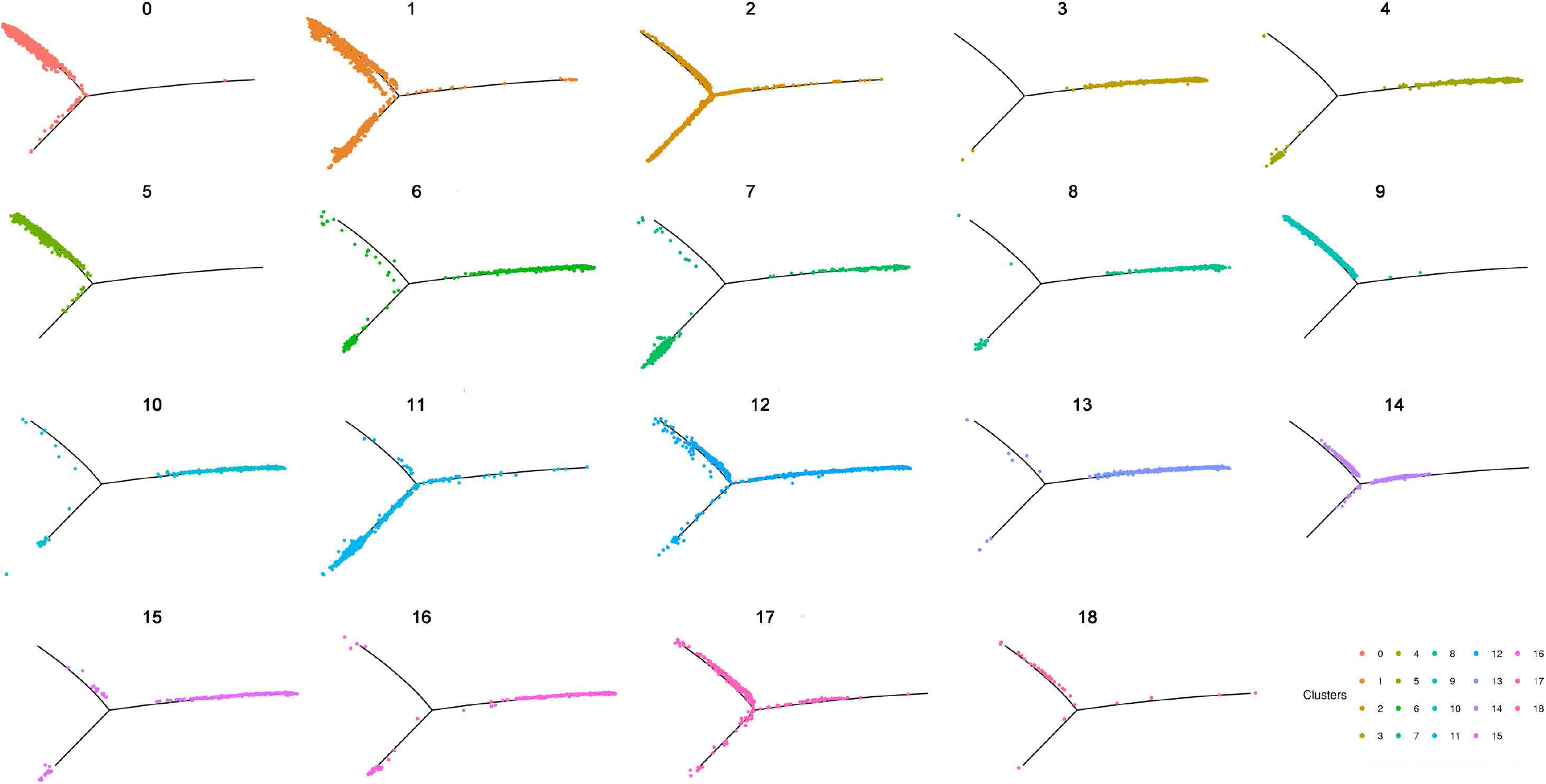
Distribution of each silkworm hemocyte cluster on pseudotime trajectories.

**Appendix Fig S3.**
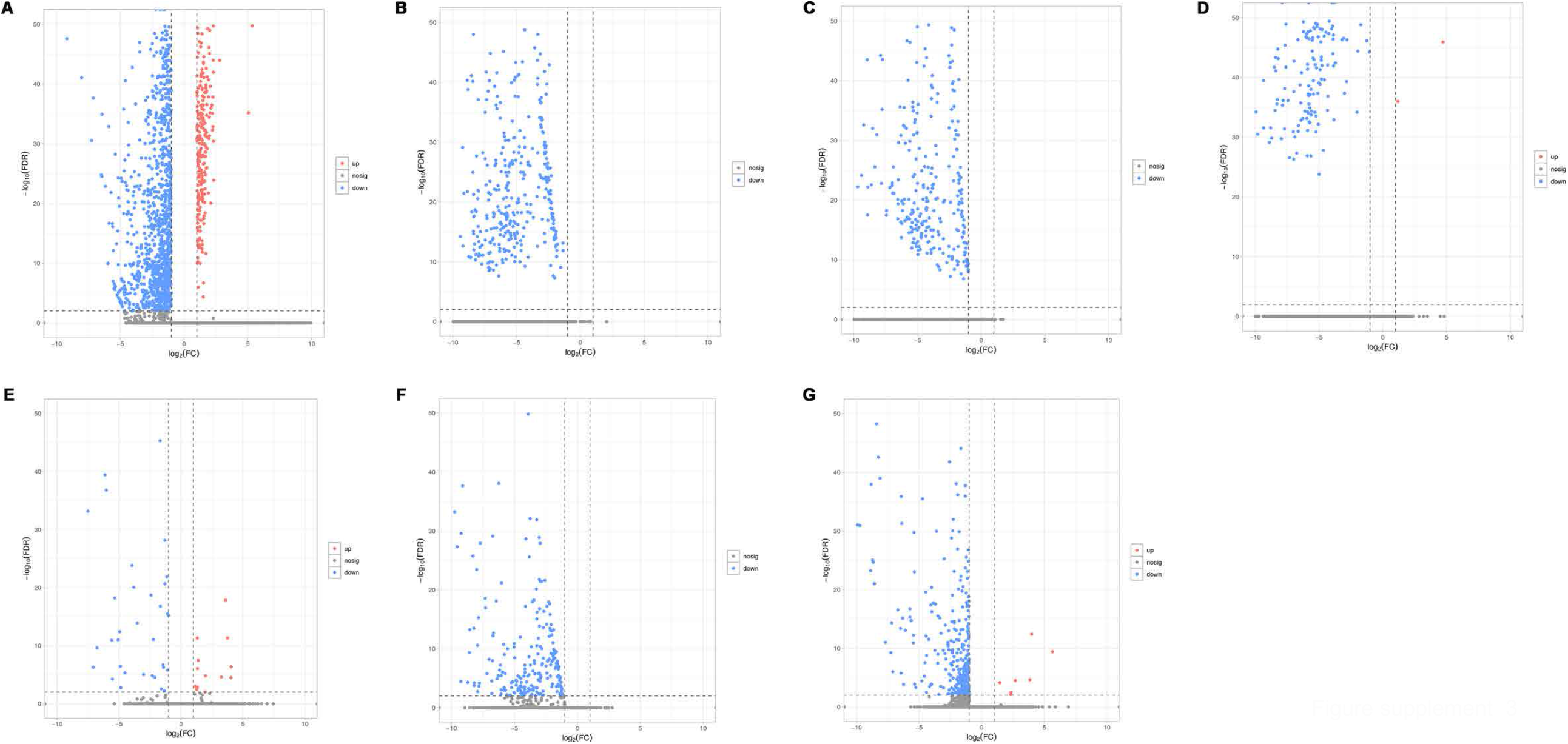
Volcano plot of identifed DEGs between BmNPV-infected and control groups in clusters 2 (A), 4 (B), 6 (C), 7 (D), 14 (E), 16 (F) and 17 (G).

**Appendix Fig S4.**
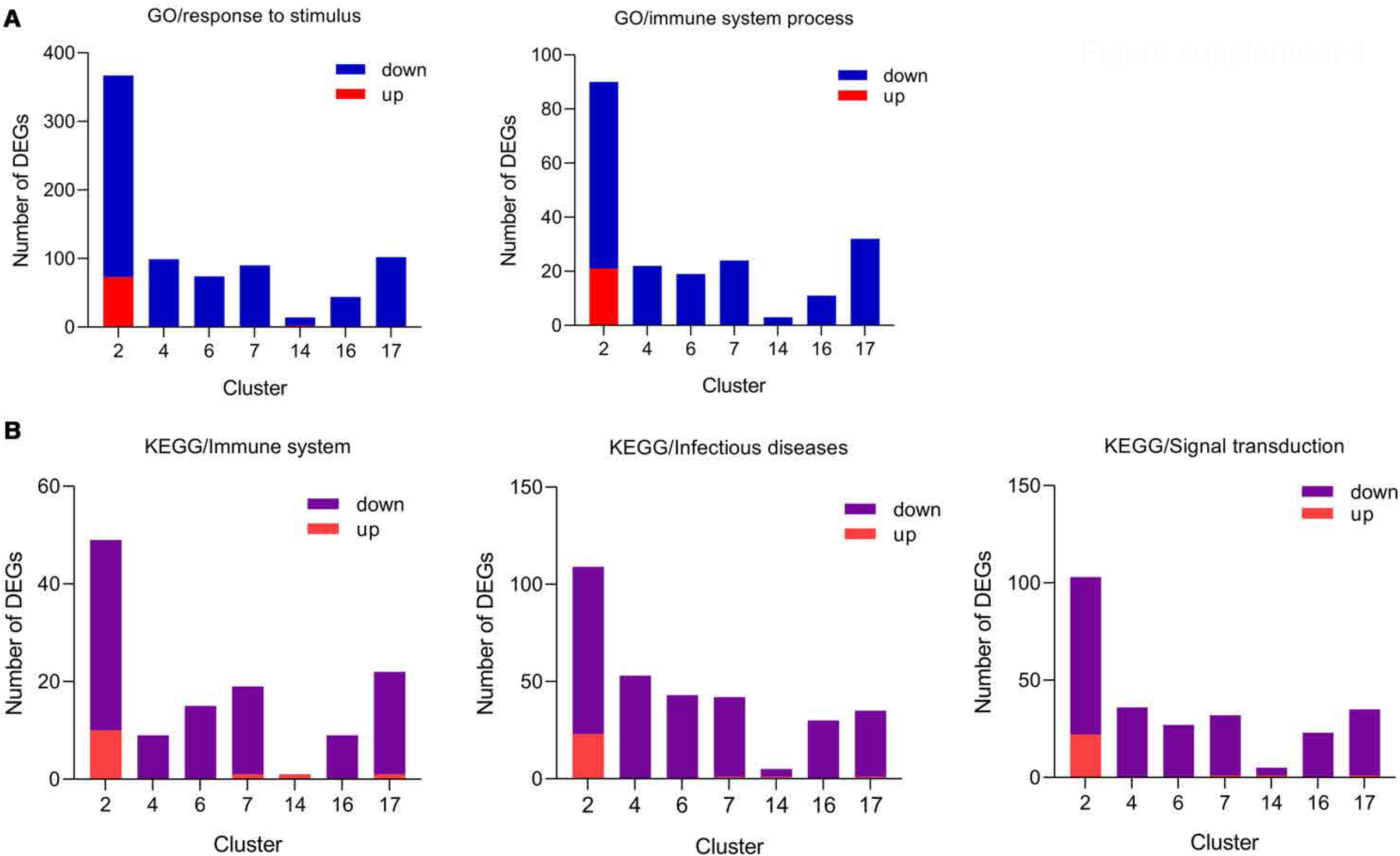
Distribution of DEGs identified between the BmNPV-infected and control groups for different viral infection-related GO terms and KEGG pathways. **(A)** DEGs involved in “immune system process” and “response to stimulus” GO terms identified in clusters 2, 4, 6, 7, 14, 16 and 17 as compared between BmNPV-infected and control cells in each cluster. **(B)** DEGs involved in “Immune system”, “Infectious diseases” and “Signal transduction” KEGG pathways identified in clusters 2, 4, 6, 7, 14, 16 and 17 as compared between BmNPV-infected and control cells in each cluster.

## References

Ahmed, S. & Kim, Y. 2019. An aquaporin mediates cell shape change required for cellular immunity in the beet armyworm, Spodoptera exigua. Scientific Reports, 9 (1):4988. doi: 10.1038/s41598-019-41541-2.

Becht, E., Mcinnes, L., Healy, J., Dutertre, C. A., Kwok, I. W. H., Ng, L. G., Ginhoux, F. & Newell, E. W. 2019. Dimensionality reduction for visualizing single-cell data using UMAP. Nature Biotechnology, 3. doi: 10.1038/nbt.4314.

Beetz, S., Brinkmann, M. & Trenczek, T. 2004. Differences between larval and pupal hemocytes of the tobacco hornworm, Manduea sexta, determined by monoclonal antibodies and density centrifugation. Journal of Insect Physiology, 50, 805–819. doi: 10.1016/j.jinsphys.2004.06.003.

Browne, N., Heelan, M. & Kavanagh, K. 2013. An analysis of the structural and functional similarities of insect hemocytes and mammalian phagocytes. Virulence, 4, 597–603. doi: 10.4161/viru.25906.

Buchon, N., Silverman, N. & Cherry, S. 2014. Immunity in Drosophila melanogaster - from microbial recognition to whole-organism physiology. Nature Reviews Immunology, 14, 796–810. doi: 10.1038/nri3763.

Cattenoz, P. B., Sakr, R., Pavlidaki, A., Delaporte, C., Riba, A., Molina, N., Hariharan, N., Mukherjee, T. & Giangrande, A. 2020. Temporal specificity and heterogeneity of Drosophila immune cells. Embo Journal, 39:e104486. doi: 10.15252/embj.2020104486.

Chen, Z. W., Nohata, J., Guo, H. Z., Li, S. L., Liu, J. Q., Guo, Y. B., Yamamoto, K., Kadono-Okuda, K., Liu, C., Arunkumar, K. P., Nagaraju, J., Zhang, Y., Liu, S. P., Labropoulou, V., Swevers, L., Tsitoura, P., Iatrou, K., Gopinathan, K. P., Goldsmith, M. R., Xia, Q. Y. & Mita, K. 2015. A comprehensive analysis of the chorion locus in silkmoth. Scientific Reports, 5:16424. doi: 10.1038/srep16424.

Chua, R. L., Lukassen, S., Trump, S., Hennig, B. P., Wendisch, D., Pott, F., Debnath, O., Thurmann, L., Kurth, F., Volker, M. T., Kazmierski, J., Timmermann, B., Twardziok, S., Schneider, S., Machleidt, F., Muller-Redetzky, H., Maier, M., Krannich, A., Schmidt, S., Balzer, F., Liebig, J., Loske, J., Suttorp, N., Eils, J., Ishaque, N., Liebert, U. G., Von Kalle, C., Hocke, A., Witzenrath, M., Goffinet, C., Drosten, C., Laudi, S., Lehmann, I., Conrad, C., Sander, L. E. & Eils, R. 2020. COVID-19 severity correlates with airway epithelium-immune cell interactions identified by single-cell analysis. Nat Biotechnol, 38, 970–979. doi: 10.1038/s41587-020-0602-4.

Clem, R. J. 2005. The role of apoptosis in defense against baculovirus infection in insects. Curr Top Microbiol Immunol, 289, 113–129. doi: 10.1007/3-540-27320-4_5.

D’Arcy, M. S. 2019. Cell death: a review of the major forms of apoptosis, necrosis and autophagy. Cell Biology International, 43, 582–592. doi: 10.1002/cbin.11137.

Davie, K., Janssens, J., Koldere, D., De Waegeneer, M., Pech, U., Kreft, L., Aibar, S., Makhzami, S., Christiaens, V., Gonzalez-Blas, C. B., Poovathingal, S., Hulselmans, G., Spanier, K. I., Moerman, T., Vanspauwen, B., Geurs, S., Voet, T., Lammertyn, J., Thienpont, B., Liu, S., Konstantinides, N., Fiers, M., Verstreken, P. & Aerts, S. 2018. A Single-Cell Transcriptome Atlas of the Aging Drosophila Brain. Cell, 174, 982-998.e20. doi: 10.1016/j.cell.2018.05.057.

Elisabeth M.M Gardiner, M. R. S. 1999. Monoclonal antibodies bind distinct classes of hemocytes in the moth Pseudoplusia includens. J Insect Physiol. (2):113–126. doi: 10.1016/s0022-1910(98)00092-4.

Feng, M., Kolliopoulou, A., Zhou, Y. H., Fei, S. G., Xia, J. M., Swevers, L. & Sun, J. C. 2020. The piRNA response to BmNPV infection in the silkworm fat body and midgut. Insect Sci. doi: 10.1111/1744-7917.12796.

Finak, G., Mcdavid, A., Yajima, M., Deng, J. Y., Gersuk, V., Shalek, A. K., Slichter, C. K., Miller, H. W., Mcelrath, M. J., Prlic, M., Linsley, P. S. & Gottardo, R. 2015. MAST: a flexible statistical framework for assessing transcriptional changes and characterizing heterogeneity in single-cell RNA sequencing data. Genome Biology, 16:278. doi: 10.1186/s13059-015-0844-5.

Fritsch, M., Gunther, S. D., Schwarzer, R., Albert, M. C., Schorn, F., Werthenbach, J. P., Schiffmann, L. M., Stair, N., Stocks, H., Seeger, J. M., Lamkanfi, M., Kronke, M., Pasparakis, M. & Kashkar, H. 2019. Caspase-8 is the molecular switch for apoptosis, necroptosis and pyroptosis. Nature, 575, 683–687. doi: 10.1038/s41586-019-1770-6.

Goldsmith, M. R., Shimada, T. & Abe, H. 2005. The genetics and genomics of the silkworm, Bombyx mori. Annual Review of Entomology, 50, 71–100. doi: 10.1146/annurev.ento.50.071803.130456.

Hartenstein, V. 2020. One too many: the surprising heterogeneity of Drosophila macrophages. Embo Journal, 39(12):e105199. doi: 10.15252/embj.2020105199.

Honti, V., Csordas, G., Kurucz, E., Markus, R. & Ando, I. 2014. The cell-mediated immunity of Drosophila melanogaster: Hemocyte lineages, immune compartments, microanatomy and regulation. Developmental and Comparative Immunology, 42, 47–56. doi: 10.1016/j.dci.2013.06.005.

Hu, X., Zhu, M., Liu, B., Liang, Z., Huang, L., Xu, J., Yu, L., Li, K., Jiang, M., Xue, R., Cao, G. & Gong, C. 2018. Circular RNA alterations in the Bombyx mori midgut following B. mori nucleopolyhedrovirus infection. Mol Immunol, 101, 461–470. doi: 10.1016/j.molimm.2018.08.008.

Hung, R. J., Hu, Y. H., Kirchner, R., Liu, Y. F., Xu, C. W., Comjean, A., Tattikota, S. G., Li, F. G., Song, W., Sui, S. H. & Perrimon, N. 2020. A cell atlas of the adult Drosophila midgut. Proceedings of the National Academy of Sciences of the United States of America, 117, 1514–1523. doi: 10.1073/pnas.1916820117.

J C GARBE, E. Y. J W FRISTROM 1993. IMP-L2:an essential secreted immunoglobulin family member implicated in neural and ectodermal development in Drosophila. Development, 119(4), 1237–1250. PMID: 8306886.

Jarvis, D. L., Bohlmeyer, D.A., Liao, Y.-F., Lomax, K.K., Merkle, R.K., Weinkauf, C., and Moremen, K.W 1997. Isolation and characterization of a class II a-mannosidase cDNA from lepidopteran insect cells. Glycobiology, 7, 113–127. doi: 10.1093/glycob/7.1.113.

Jevitt, A., Chatterjee, D., Xie, G. Q., Wang, X. F., Otwell, T., Huang, Y. C. & Deng, W. M. 2020. A single-cell atlas of adult Drosophila ovary identifies transcriptional programs and somatic cell lineage regulating oogenesis. Plos Biology, 18 (4):e3000538. doi: 10.1371/journal.pbio.3000538.

Kim, D., Landmead, B. & Salzberg, S. L. 2015. HISAT: a fast spliced aligner with low memory requirements. Nature Methods, 12, 357–360. doi: 10.1038/nmeth.3317.

Kingsolver, M. B., Huang, Z. J. & Hardy, R. W. 2013. Insect Antiviral Innate Immunity: Pathways, Effectors, and Connections. Journal of Molecular Biology, 425, 4921–4936. doi: 10.1016/j.jmb.2013.10.006.

Koczka, K., Peters, P., Ernst, W., Himmelbauer, H., Nika, L. & Grabherr, R. 2018. Comparative transcriptome analysis of a Trichoplusia ni cell line reveals distinct host responses to intracellular and secreted protein products expressed by recombinant baculoviruses. Journal of Biotechnology, 270, 61–69. doi: 10.1016/j.jbiotec.2018.02.001.

Kolliopoulou, A., Van Nieuwerburgh, F., Stravopodis, D. J., Deforce, D., Swevers, L. & Smagghe, G. 2015. Transcriptome Analysis of Bombyx mori Larval Midgut during Persistent and Pathogenic Cytoplasmic Polyhedrosis Virus Infection. Plos One, 10 (3):e0121447. doi: 10.1371/journal.pone.0121447.

Krejcova, G., Danielova, A., Nedbalova, P., Kazek, M., Strych, L., Chawla, G., Tennessen, J. M., Lieskovska, J., Jindra, M., Dolezal, T. & Bajgar, A. 2019.Drosophila macrophages switch to aerobic glycolysis to mount effective antibacterial defense. Elife, 8: e50414. doi: 10.7554/eLife.50414.

Lavine, M. D. & Strand, M. R. 2002. Insect hemocytes and their role in immunity. Insect Biochemistry and Molecular Biology, 32, 1295–1309. doi: 10.1016/s0965-1748(02)00092-9.

Li, G., Qian, H., Luo, X., Xu, P., Yang, J., Liu, M. & Xu, A. 2016. Transcriptomic Analysis of Resistant and Susceptible Bombyx mori Strains Following BmNPV Infection Provides Insights into the Antiviral Mechanisms. Int J Genomics, 2016, 2086346. doi: 10.1155/2016/2086346.

Linderman, G. C., Rachh, M., Hoskins, J. G., Steinerberger, S. & Kluger, Y. 2019. Fast interpolation-based t-SNE for improved visualization of single-cell RNA-seq data. Nature Methods, 16, 243– 245. doi: 10.1038/s41592-018-0308-4.

Liu, F., Xu, Q., Zhang, Q., Lu, A., Beerntsen, B. T. & Ling, E. 2013. Hemocytes and hematopoiesis in the silkworm, Bombyx mori. Isj-Invertebrate Survival Journal, 10, 102–109.

Liu, W. T., Tu, W. C., Lin, C. H., Yang, U. C. & Chen, C. C. 2017. Involvement of cecropin B in the formation of the Aedes aegypti mosquito cuticle. Scientific Reports, 7 (1):16395. doi: 10.1038/s41598-017-16625-6.

Lu, F., Wei, Z., Luo, Y., Guo, H., Zhang, G., Xia, Q. & Wang, Y. 2020. SilkDB 3.0: visualizing and exploring multiple levels of data for silkworm. Nucleic Acids Res, 48, D749–D755. doi: 10.1093/nar/gkz919.

Marques, J. T. & Imler, J. L. 2016. The diversity of insect antiviral immunity: insights from viruses. Current Opinion in Microbiology, 32, 71–76. doi: 10.1016/j.mib.2016.05.002.

Matsumoto, H., Tsuzuki, S., Date-Ito, A., Ohnishi, A. & Hayakawa, Y. 2012. Characteristics common to a cytokine family spanning five orders of insects. Insect Biochemistry and Molecular Biology, 42, 446–454. doi: 10.1016/j.ibmb.2012.03.001.

Mcdavid, A., Finak, G., Chattopadyay, P. K., Dominguez, M., Lamoreaux, L., Ma, S. S., Roederer, M. & Gottardo, R. 2013. Data exploration, quality control and testing in single-cell qPCR-based gene expression experiments. Bioinformatics, 29, 461–467. doi: 10.1093/bioinformatics/bts714.

Nakahara, Y., Kanamori, Y., Kiuchi, M. & Kamimura, M. 2010. Two Hemocyte Lineages Exist in Silkworm Larval Hematopoietic Organ. Plos One, 5 (7):e11816. doi: 10.1371/journal.pone.0011816.

Nakahara, Y., Shimura, S., Ueno, C., Kanamori, Y., Mita, K., Kiuchi, M. & Kamimura, M. 2009. Purification and characterization of silkworm hemocytes by flow cytometry. Dev Comp Immunol, 33, 439–448. doi: 10.1016/j.dci.2008.09.005.

Nguyen, Q., Chan, L. C. L., Nielsen, L. K. & Reid, S. 2013a. Genome scale analysis of differential mRNA expression of Helicoverpa zea insect cells infected with a H. armigera baculovirus. Virology, 444, 158–170. doi: 10.1016/j.virol.2013.06.004.

Nguyen, Q., Nielsen, L. K. & Reid, S. 2013b. Genome Scale Transcriptomics of Baculovirus-Insect Interactions. Viruses-Basel, 5, 2721–2747. doi: 10.3390/v5112721.

Pertea, M., Kim, D., Pertea, G. M., Leek, J. T. & Salzberg, S. L. 2016. Transcript-level expression analysis of RNA-seq experiments with HISAT, StringTie and Ballgown. Nature Protocols, 11, 1650–1667. doi: 10.1038/nprot.2016.095.

Ramond, E., Dudzic, J. P. & Lemaitre, B. 2020. Comparative RNA-Seq analyses of Drosophila plasmatocytes reveal gene specific signatures in response to clean injury and septic injury. Plos One, 15 (6):e0235294. doi: 10.1371/journal.pone.0235294.

Ribeiro, C. & Brehelin, M. 2006. Insect haemocytes: What type of cell is that? Journal of Insect Physiology, 52, 417–429. doi: 10.1016/j.jinsphys.2006.01.005.

Rohrmann, G. F. 2019. In: TH (ed.) Baculovirus Molecular Biology. Bethesda (MD). Sarraf-Zadeh, L., Christen, S., Sauer, U., Cognigni, P., Miguel-Aliaga, I., Stocker, H., Kohler, K. & Hafen, E. 2013. Local requirement of the Drosophila insulin binding protein imp-L2 in coordinating developmental progression with nutritional conditions. Developmental Biology, 381, 97–106. doi: 10.1016/j.ydbio.2013.06.008.

Satija, R., Farrell, J. A., Gennert, D., Schier, A. F. & Regev, A. 2015. Spatial reconstruction of single-cell gene expression data. Nature Biotechnology, 33, 495–502. doi: 10.1038/nbt.3192.

Sheehan, G., Farrell, G. & Kavanagh, K. 2020. Immune priming: the secret weapon of the insect world. Virulence, 11, 238–246. doi: 10.1080/21505594.2020.1731137.

Shi, J. J., Gao, W. Q. & Shao, F. 2017. Pyroptosis: Gasdermin-Mediated Programmed Necrotic Cell Death. Trends in Biochemical Sciences, 42, 245–254. doi: 10.1016/j.tibs.2016.10.004.

Slaidina, M., Banisch, T. U., Gupta, S. & Lehmann, R. 2020. A single-cell atlas of the developing Drosophila ovary identifies follicle stem cell progenitors. Genes & Development, 34, 239–249. doi: 10.1101/gad.330464.119.

Steuerman, Y., Cohen, M., Peshes-Yaloz, N., Valadarsky, L., Cohn, O., David, E., Frishberg, A., Mayo, L., Bacharach, E., Amit, I. & Gat-Viks, I. 2018. Dissection of Influenza Infection In Vivo by Single-Cell RNA Sequencing. Cell Systems, 6, 679-691.e4. doi: 10.1016/j.cels.2018.05.008.

Strand, M. R. 2008. The insect cellular immune response. Insect Science, 15, 1–14. doi: 10.1111/j.1744-7917.2008.00183.x.

Stuart, T., Butler, A., Hoffman, P., Hafemeister, C., Papalexi, E., Mauck, W. M., Hao, Y. H., Stoeckius, M., Smibert, P. & Satija, R. 2019. Comprehensive Integration of Single-Cell Data. Cell, 177, 1888-1902.e21. doi: 10.1016/j.cell.2019.05.031.

Swevers, L., Liu, J. & Smagghe, G. 2018. Defense Mechanisms against Viral Infection in Drosophila: RNAi and Non-RNAi. Viruses-Basel, 10 (5):230. doi: 10.3390/v10050230.

Tanaka, H., Ishibashi, J., Fujita, K., Nakajima, Y., Sagisaka, A., Tomimoto, K., Suzuki, N., Yoshiyama, M., Kaneko, Y., Iwasaki, T., Sunagawa, T., Yamaji, K., Asaoka, A., Mita, K. & Yamakawa, M. 2008. A genome-wide analysis of genes and gene families involved in innate immunity of Bombyx mori. Insect Biochemistry and Molecular Biology, 38, 1087–1110. doi: 10.1016/j.ibmb.2008.09.001.

Tattikota, S. G., Cho, B., Liu, Y. F., Hu, Y. H., Barrera, V., Steinbaugh, M. J., Yoon, S. H., Comjean, A., Li, F. G., Dervis, F., Hung, R. J., Nam, J. W., Sui, S. H., Shim, J. & Perrimon, N. 2020. A single-cell survey of Drosophila blood. Elife, 9: e54818. doi: 10.7554/eLife.54818.

Trapnell, C., Cacchiarelli, D., Grimsby, J., Pokharel, P., Li, S. Q., Morse, M., Lennon, N. J., Livak, K. J., Mikkelsen, T. S. & Rinn, J. L. 2014. The dynamics and regulators of cell fate decisions are revealed by pseudotemporal ordering of single cells. Nature Biotechnology, 32, 381 –386. doi: 10.1038/nbt.2859.

Wilk, A. J., Rustagi, A., Zhao, N. Q., Roque, J., Martinez-Colon, G. J., Mckechnie, J. L., Ivison, G. T., Ranganath, T., Vergara, R., Hollis, T., Simpson, L. J., Grant, P., Subramanian, A., Rogers, A. J. & Blish, C. A. 2020. A single-cell atlas of the peripheral immune response in patients with severe COVID-19. Nat Med, 26, 1070–1076. doi: 10.1038/s41591-020-0944-y.

Xia, Q., Li, S. & Feng, Q. 2014. Advances in silkworm studies accelerated by the genome sequencing of Bombyx mori. Annu Rev Entomol, 59, 513–536. doi: 10.1146/annurev-ento-011613-161940.

Y Kato, K. T. H Hirochika, M Yamakawa 1993. Expression and characterization of cDNAs for cecropin B, an antibacterial protein of the silkworm, Bombyx mori. Insect Biochem Mol Biol, 23(2), 285–290. doi: 10.1016/0965-1748(93)90009-h.

Yamashita, M. & Iwabuchi, K. 2001. Bombyx mori prohemocyte division and differentiation in individual microcultures. J Insect Physiol, 47, 325–331. doi: 10.1016/s0022-1910(00)00144-x.

Yu, H., Wang, X., Xu, J., Ma, Y., Zhang, S., Yu, D., Fei, D. & Muhammad, A. 2017. iTRAQ-based quantitative proteomics analysis of molecular mechanisms associated with Bombyx mori (Lepidoptera) larval midgut response to BmNPV in susceptible and near-isogenic strains. J Proteomics, 165, 35–50. doi: 10.1016/j.jprot.2017.06.007.

Zhang, J., Liu, J., Yuan, Y., Huang, F., Ma, R., Luo, B., Xi, Z., Pan, T., Liu, B., Zhang, Y., Zhang, X., Luo, Y., Wang, J., Zhao, M., Lu, G., Deng, K. & Zhang, H. 2020a. Twowaves of pro-inflammatory factors are released during the influenza A virus (IAV)-driven pulmonary immunopathogenesis. PLoS Pathog, 16, e1008334. doi: 10.1371/journal.ppat.1008334.

Zhang, K., Li, C., Weng, X., Su, J., Shen, L., Pan, G., Long, D., Zhao, A. & Cui, H.2018. Transgenic characterization of two silkworm tissue-specific promoters in the haemocyte plasmatocyte cells. Insect Mol Biol, 27, 133–142. doi: 10.1111/imb.12360.

Zhang, K., Tan, J., Xu, M., Su, J., Hu, R., Chen, Y., Xuan, F., Yang, R. & Cui, H. 2014. A novel granulocyte-specific alpha integrin is essential for cellular immunity in the silkworm Bombyx mori. J Insect Physiol, 71, 61–67. doi: 10.1016/j.jinsphys.2014.10.007.

Zhang, S., Yin, H., Shen, M., Huang, H., Hou, Q., Zhang, Z., Zhao, W., Guo, X. & Wu, P. 2020b. Analysis of lncRNA-mediated gene regulatory network of Bombyx mori in response to BmNPV infection. J Invertebr Pathol, 170, 107323. doi: 10.1016/j.jip.2020.107323.

Zhang, S. Z., Wang, J., Zhu, L. B., Toufeeq, S., Xu, X., You, L. L., Li, B., Hu, P. & XU,J. P. 2020c. Quantitative label-free proteomic analysis reveals differentially expressed proteins in the digestive juice of resistant versus susceptible silkworm strains and their predicted impacts on BmNPV infection. J Proteomics, 210, 103527. doi: 10.1016/j.jprot.2019.103527.

Zhu, L., Yang, P., Zhao, Y., Zhuang, Z., Wang, Z., Song, R., Zhang, J., Liu, C., Gao, Q., Xu, Q., Wei, X., Sun, H. X., Ye, B., Wu, Y., Zhang, N., Lei, G., Yu, L., Yan, J., Diao, G., Meng, F., Bai, C., Mao, P., Yu, Y., Wang, M., Yuan, Y., Deng, Q., Li, Z., Huang, Y.,Hu, G., Liu, Y., Wang, X., Xu, Z., Liu, P., Bi, Y., Shi, Y., Zhang, S., Chen, Z., Wang, J., Xu, X., Wu, G., Wang, F. S., Gao, G. F., Liu, L. & Liu, W. J. 2020. Single-Cell Sequencing of Peripheral Mononuclear Cells Reveals Distinct Immune Response Landscapes of COVID-19 and Influenza Patients. Immunity, 53(3):685-696.e3. doi: 10.1016/j.immuni.2020.07.009.

Zimmermann, L., Stephens, A., Nam, S. Z., Rau, D., Kubler, J., Lozajic, M., Gabler, F., Soding, J., Lupas, A. N. & Alva, V. 2018. A Completely Reimplemented MPI Bioinformatics Toolkit with a New HHpred Server at its Core. Journal of Molecular Biology, 430, 2237–2243. doi: 10.1016/j.jmb.2017.12.007.

